# IL-21 Shapes the B Cell Response in a Context-Dependent Manner

**DOI:** 10.1101/2024.07.13.600808

**Authors:** Youngjun Kim, Francesca Manara, Simon Grassmann, Kalina T. Belcheva, Kanelly Reyes, Hyunu Kim, Stephanie Downs-Canner, William T. Yewdell, Joseph C. Sun, Jayanta Chaudhuri

## Abstract

The cytokine interleukin-21 (IL-21) is a pivotal T cell-derived signal crucial for germinal center (GC) responses, but the precise mechanisms by which IL-21 influences B cell function remain elusive. Here, we investigated the B cell-intrinsic role of IL-21 signaling by employing a novel IL-21 receptor (*Il21r*) conditional knock-out mouse model and *ex vivo* culture systems and uncovered a surprising duality of IL-21 signaling in B cells. While IL-21 stimulation of naïve B cells led to Bim-dependent apoptosis, it promoted robust proliferation of pre-activated B cells, particularly class-switched IgG1^+^ B cells *ex vivo*. Consistent with this, B cell-specific deletion of *Il21r* led to a severe defect in IgG1 responses *in vivo* following immunization. Intriguingly, *Il21r*-deleted B cells are significantly impaired in their ability to transition from a pre-GC to a GC state following immunization. Although *Il21r*-deficiency did not affect the proportion of IgG1^+^ B cells among GC B cells, it greatly diminished the proportion of IgG1^+^ B cells among the plasmablast/plasma cell population. Collectively, our data suggest that IL-21 serves as a critical regulator of B cell fates, influencing B cell apoptosis and proliferation in a context-dependent manner.

## INTRODUCTION

During a T cell-dependent immune response, B cells undergo a sophisticated orchestration of events, beginning with their activation and migration to the border between the T cell zone and B cell follicles^1^. Here, they engage in interactions with cognate T cells and initiate activation-induced cytidine deaminase (AID)-driven immunoglobulin heavy chain (IgH) class switch recombination (CSR)^2,3^, a DNA deletional recombination event that changes the isotype of the expressed antibody from IgM to a secondary isotype such as IgG, IgE, or IgA. Subsequently, a subset of activated B cells translocate to the center of the follicles, where they mature into germinal center (GC) B cells^4–6^.

GCs are microanatomical structures comprised of two major compartments – dark zone (DZ) and light zone (LZ). In the DZ, B cells undergo robust proliferation and AID-mediated somatic hypermutation (SHM), generating B cell receptors (BCRs) with varying antigen-affinity. The DZ B cells subsequently migrate into the LZ, where their newly minted BCRs test their affinity for antigens bound to follicular dendritic cells (FDCs). Clones with higher BCR affinities are more likely to capture, internalize, and present antigens to follicular helper T (T_FH_) cells. Since GC B cells are poised to undergo apoptosis, and require signals from T_FH_ cells for positive selection, high-affinity clones are more likely to be preferentially selected than low-affinity clones^7–9^. Positively selected B cell clones upregulate c-Myc, initiate cell cycle, and transition into the DZ^10,11^. This iterative cycling between the DZ and LZ enforces rigorous selection for high-affinity B cell clones, akin to a process of Darwinian selection, contributing to antibody affinity maturation within GCs.

IL-21, a pleiotropic cytokine primarily produced by T_FH_ cells, natural killer T (NKT) cells, and T_H_17 cells, signals through a heterodimeric receptor composed of the IL-21 receptor (IL-21R) subunit and the common gamma chain (γc)^12,13^. IL-21R is expressed on a variety of hematopoietic cells including dendritic cells, macrophages, NK cells, and on B cells^13^. Upon binding to the receptor, IL-21 triggers activation of downstream signaling pathways mediated by STAT1, STAT3, STAT5, PI3K-Akt, and MAPK^12,14–16^. Studies employing IL-21 or IL-21R-deficient mice have demonstrated the role of IL-21 signaling in sustaining both GC B cells and T_FH_ cells, profoundly impacting antibody and GC responses^17–20^. It has been suggested that IL-21 acts on B cells to promote GC formation, plasma cell (PC) differentiation, and memory B cell responses^19–22^. These studies, however, predominantly relied on *Il21^-/-^* or *Il21r^-/-^* mouse models, which abrogate IL-21R signaling across all cell populations, including T_FH_ cells, thereby limiting insights into the B-cell intrinsic role of IL-21.

Here, we developed a novel conditional *Il21r* knock-out mouse model to facilitate an extensive examination of the B cell-intrinsic role of IL-21 signaling. Using both *ex vivo* primary B cell cultures and *in vivo* immunization models, we uncovered a context-dependent influence of IL-21 signaling on B cell fate. We found that while IL-21 induces apoptosis in naïve B cells, it supports proliferation in activated B cells *ex vivo*. Notably, IL-21 treatment led to a selective expansion of the class-switched B cells, likely due to cooperative effects with BCR signaling. Furthermore, B cell-specific deletion of *Il21r* in mice led to a marked defect in pre-GC to GC transition, accompanied by impaired affinity maturation, alterations in the composition of light and dark zones, and a stricter selection for antigen-specific B cells. Additionally, IL-21 deficiency leads to impaired plasmablast (PB)/plasma cell (PC) differentiation and a dramatic reduction in IgG1^+^ cells within this population. Our results suggest that IL-21 functions as a dynamic regulator of B cell fates during T-dependent responses, critically influencing the dynamics of GC responses.

## RESULTS

### IL-21 induces apoptosis in naïve B cells

To investigate the role of IL-21 in B cell activation and proliferation during a T-dependent response, we cultured naïve splenic B cells from wild-type C57BL/6 mice with cytokine cocktails designed to simulate interactions of B cells with cognate T cells: anti-CD40 antibody (αCD40) alone, αCD40 + IL-4, αCD40 + IL-21, or αCD40 + IL-4 + IL-21 (**Figure 1a**). While αCD40-stimulated B cell numbers remained largely unchanged over time, supplementing the culture with IL-4 resulted in a robust proliferation. Surprisingly, addition of IL-21 to either αCD40 or αCD40 + IL-4 stimulation resulted in a significant reduction in cell number (**Figure 1b**) and a marked increase in the frequency of active caspase-3^+^ cells, indicative of apoptosis (**Figure 1c**). This trend persisted even at a five-fold lower concentration of IL-21 (10 ng/mL) (**Supplementary Figure 1a, b**).

**Figure 1.**
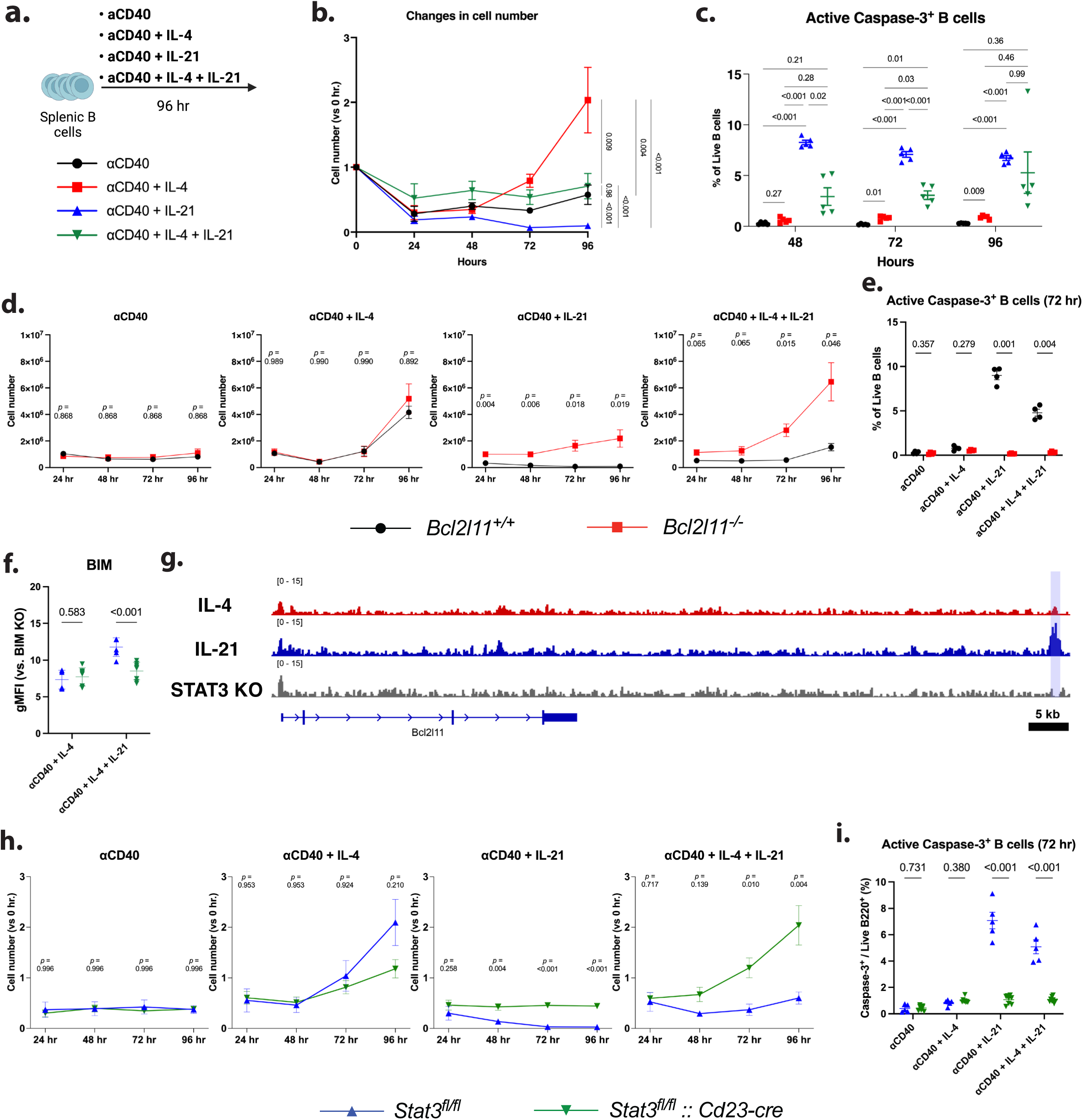
IL-21 promotes apoptosis of naïve B cells through STAT3-Bim axis *ex vivo*. (a) Illustration of activation protocol of naïve primary B cells *ex vivo*. Naïve splenic B cells were cultured with αCD40 (1 μg/mL), IL-4 (12.5 ng/mL), or IL-21 (50 ng/mL). (b) Changes in cell numbers in culture over time. The y-axis represents changes in cell numbers relative to the starting point (0 hours). (c) Percentage of active caspase-3^+^ B cells among total live B cells. (d, h) Changes in cell numbers over time under various stimulation conditions. The y-axis represents the (d) absolute cell number or (h) changes in cell numbers relative to the starting point (0 hours). (e, i) Percentage of active caspase-3^+^ B cells among the total live B cell population in different stimulation conditions at 72 hours after starting culture. (f) Geometric MFI of Bim in STAT3-deficient or control B cells stimulated as indicated for 8 hours. (g) Splenic B cells were pre-stimulated with αCD40 + IL-4 for 48 hours and FACS-sorted IgM+ B cells were re-stimulated with either αCD40 + IL-4 or αCD40 + IL-21 for 30 minutes. Chromatin binding of STAT3 was subsequently assessed by CUT&RUN and spike-in scaled signals were averaged for track visualization. Blue highlight denotes a significant peak called by MACS2 algorithm for pooled samples of WT IL-21 condition. Data were pooled from two or three independent experiments with *n* = 2 or 3 per experiment. (a-c) Different stimulation conditions are represented by distinct colors: αCD40 (black), αCD40 + IL-4 (red), αCD40 + IL-21 (blue), and αCD40 + IL-4 + IL-21 (green). (d-h) Different experimental groups are represented by distinct colors: *Bcl2l11^+/+^* (black), *Bcl2l11^-/-^* (red), *Stat3^fl/fl^* (blue), and *Stat3^fl/fl^::Cd23-cre* (green). For (b), the 96-hour timepoint data were subject to ANOVA in conjunction with Tukey’s multiple comparisons test. Data in (c) were analyzed with two-way ANOVA in conjunction with Tukey’s multiple comparisons test. Data in (d), (e), (f), (h), and (i) were analyzed with Student’s unpaired t-test, with multiple comparisons correction applied using the Holm-Šídák method.

To determine the signaling pathways underlying IL-21-induced B cell apoptosis, we performed RNA-sequencing (RNA-seq) on naïve B cells stimulated *ex vivo* with combinations of αCD40, IL-4, and IL-21 for 6 or 24 hours (**Supplementary Figure 2a**). Principal component analysis (PCA) illustrated clustering of samples based on timepoint and stimulation (**Supplementary Figure 2b**). As expected, IL-21 stimulation led to enrichment of various signaling pathways including the MAPK and PI3K-Akt pathways^16^. We also observed downregulation of pathways involved in DNA replication, ribosome biogenesis, and oxidative phosphorylation upon IL-21 stimulation, suggesting impaired cellular fitness (**Supplementary Figure 2c**). Strikingly, IL-21 stimulation downregulated anti-apoptotic genes (e.g., *Bcl2, Bcl2l1*) and upregulated pro-apoptotic genes (e.g., *Pmaip1, Bcl2l11*) (**Supplementary Figure 2d-g**). Collectively, these data suggest that IL-21 promotes apoptosis in naïve B cells *ex vivo*.

### IL-21 drives apoptosis of B cells through the STAT3-Bim axis

IL-21 stimulation induced upregulation of *Bcl2l11*, the gene encoding the pro-apoptotic factor Bim (**Supplementary Figure 2d, g**). To examine if IL-21 induces apoptosis through a Bim-dependent pathway, we cultured Bim-deficient splenic B cells with αCD40, αCD40 + IL-4, αCD40 + IL-21, or αCD40 + IL-4 + IL-21. The number of Bim-deficient B cells was comparable to that of Bim-sufficient WT B cells in the absence of IL-21. However, in the presence of IL-21, the absolute number of Bim-deficient B cells was significantly higher than WT B cells (**Figure 1d**). Additionally, Bim deficiency reduced the frequency of active caspase-3^+^ cells to virtually zero upon IL-21 stimulation (**Figure 1e**).

Given that STAT3 acts as the major downstream signal transducer of IL-21^13^, we hypothesized that STAT3 upregulates Bim upon IL-21 stimulation in B cells. To test this, we bred *Stat3^fl/fl^* mice with *Cd23-cre* mice to delete STAT3 in mature B cells. We observed that in B cells exposed to IL-21, STAT3-deficiency significantly reduced Bim expression (**Figure 1f)**. Additionally, we carried out CUT&RUN analysis to examine chromatin localization of STAT3 and observed a significant STAT3 peak at the *Bcl2l11* locus in IL-21-stimulated B cells, suggesting that STAT3 directly regulates Bim expression (**Figure 1g)**. Finally, we observed that STAT3-deficient B cells were protected from IL-21 mediated cell attrition and apoptosis similar to Bim-deficient B cells (**Figure 1h, i**). Collectively, these data suggest that IL-21 promotes B cell apoptosis through the STAT3-Bim axis.

Consistent with these notions, transient overexpression of IL-21 in mice using hydrodynamic injection^23^ of *Il21*-encoding plasmids (**Supplementary Figure 3a)** led to increased levels of phosphorylated STAT3 in naïve splenic B cells (**Supplementary Figure 3b)**. This was accompanied by a reduction in mature follicular (Fo) and marginal zone (MZ) B cell populations, coupled with an increase in transitional 1 (T1) B cells, suggesting increased turnover of the mature B cell pool in IL-21 overexpressed animals (**Supplementary Figure 3c, d)**. Interestingly, IL-21 overexpression led to a significant increase in IL-21R surface expression across all splenic B cell subsets (**Supplementary Figure 3e)**, consistent with increased *Il21r* transcription observed in B cells supplemented with IL-21 *ex vivo* (**Supplementary Figure 2i)**. This upregulation suggests a potential positive feedback loop, wherein IL-21 signaling enhances the expression of its own receptor.

### Strong BCR signaling protects B cells from IL-21 mediated cell attrition *ex vivo*

*In vivo*, B cells predominantly encounter cognate T cell help at two key locations: the B-T border and the LZ of GCs, where BCR signaling plays a critical role^24^. Upon antigen engagement, B cells upregulate CCR7 expression, facilitating their localization at the B-T border^1^. Additionally, BCR signaling promotes positive selection in GCs^25^. Given the significance of BCR signaling in B cell activation and GC responses, we investigated whether BCR signaling influences B cell responses to IL-21 stimulation by culturing naïve splenic WT B cells with combinations of αCD40, IL-4, and IL-21, in the presence or absence of BCR-crosslinking using anti-IgM monoclonal antibody (αIgM). Notably, BCR crosslinking led to a significant increase in cell numbers across all four stimulation conditions (**Figure 2a**). Furthermore, αIgM treatment significantly reduced the proportion of active caspase-3^+^ population in αCD40 + IL-21 stimulated B cells (**Figure 2b**), particularly at the 48-hour time point. These data suggest that BCR signaling protects B cells from IL-21-mediated apoptosis.

**Figure 2.**
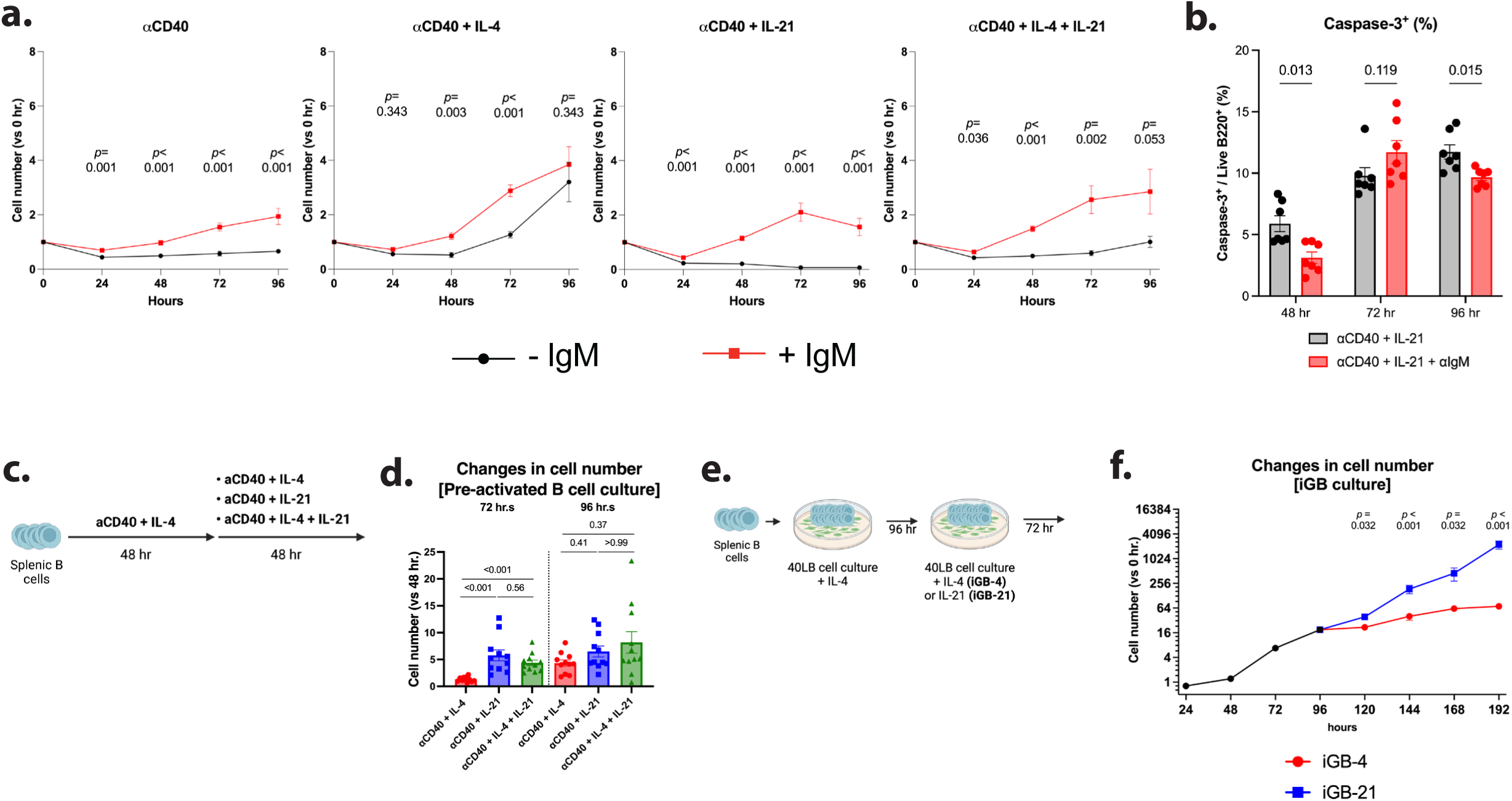
BCR cross-linking or pre-activation of B cells mitigate IL-21-dependent apoptosis. (a) Changes in cell numbers over time under various stimulation conditions. (b) Percentage of active caspase-3^+^ B cells among the total live B cells. (c) Schematic representation of pre-activated B cell culture. Naïve splenic B cells were cultured with αCD40 + IL-4 for 48 hours, followed by secondary stimulation with αCD40 + IL-4, αCD40 + IL-21, or αCD40 + IL-4 + IL-21 for an additional 48 hours. Concentration for stimuli: αCD40 (1 μg/mL), IL-4 (12.5 ng/mL), IL-21 (50 ng/mL). (d) Changes in cell numbers at 72 and 96 hours of culture compared to the 48-hour time point. The y-axis represents changes in cell numbers relative to the beginning of secondary stimulation (48 hour). (e) Schematic depiction of the iGB culture setup. Naïve splenic B cells were cultured with the 40LB feeder cells and IL-4 (1 ng/mL) for 96 hours. Subsequently, secondary stimulation was performed with fresh 40LB feeder cells in the presence of IL-4 (1 ng/mL) or IL-21 (10 ng/mL). (f) Changes in cell numbers over time. The y-axis represents changes in cell numbers relative to the start of cell culture (0 hours). Data were pooled from three independent experiments with *n* = 3 or 4 per experiment. Data in (a), (b), and (f) were analyzed with Student’s unpaired t-test, with multiple comparisons correction applied using the Holm-Šídák method. Data in (d) were statistically analyzed using ANOVA in conjunction with Tukey’s multiple comparisons test.

### IL-21 drives proliferation in activated B cells

Given that GC B cells are primed by various T cell-derived stimuli before reaching the GCs, where they interact with T_FH_ cells, we investigated the impact of IL-21 stimulation on pre-activated B cells. We initially exposed B cells to αCD40 + IL-4 for 48 hours, followed by additional stimulation with αCD40 + IL-4, αCD40 + IL-21, or αCD40 + IL-4 + IL-21 for another 48 hours (**Figure 2c**). In stark contrast to the death observed when naïve B cells were continuously exposed to IL-21 (**Figure 1b, c**), the addition of IL-21 to pre-activated B cells triggered a significant increase in the cell number within 24 hours (72 hours after the start of culture) compared to cells not supplemented with IL-21 (**Figure 2d**). However, this burst was not sustained at 48 hours (96 hours after the start of culture) (**Figure 2d**), likely due to the typical decline in proliferative capability and viability observed around 96 hours of *ex vivo* culture.

To circumvent this limitation, we adopted the induced GC B cell (iGB) culture system that allows culturing B cells for at least 8 days without losing proliferative capacity^26^. In this system, splenic B cells are co-cultured with 40LB feeder cells expressing CD40 ligand (CD40L) and B-cell activating factor (BAFF) in the presence of IL-4 for 96 hours. Subsequently, cells are either exposed to IL-21 (iGB-21) or maintained in culture with IL-4 (iGB-4) (**Figure 2e**). Remarkably, iGB-21 cells exhibited more robust cell proliferation than iGB-4 at all time points (**Figure 2f**). Therefore, unlike naïve B cells, pre-activated B cells undergo significant expansion upon IL-21 stimulation.

### IL-21 drives expansion of IgG1^+^ B cells without inducing CSR to IgG1 *ex vivo*

During an immune response, B cells undergo CSR to produce antibodies tailored to combat specific pathogens. Different classes of pathogens create distinct immune microenvironments, prompting CSR to different antibody isotypes. We investigated the role of IL-21 in CSR by culturing naïve B cells with αCD40, αCD40 + IL-4, αCD40 + IL-21, or αCD40 + IL-4 + IL-21. Since both IL-4 and IL-21 are known to influence IgG1 responses^17,27–29^, our analysis focused on CSR to IgG1. While αCD40 alone induced less than 1% of class switching to IgG1, the addition of IL-21 resulted in 25% of surviving B cells expressing IgG1. Stimulation with αCD40 + IL-4 induced robust CSR to IgG1, with approximately 30% of B cells being IgG1^+^ at 96 hours. Notably, addition of IL-21 on top of αCD40 + IL-4 led to a two-fold increase in the percentage of IgG1^+^ B cells (**Figure 3a, b)**. These findings were consistent even at a five-fold lower concentration of IL-21 (**Supplementary Figure 1c**). Additionally, IL-21 drove an increase in the proportion of IgG1^+^ B cells in both pre-activated B cells (**Figure 3c, d)** and iGB cells (**Figure 3e)**.

**Figure 3.**
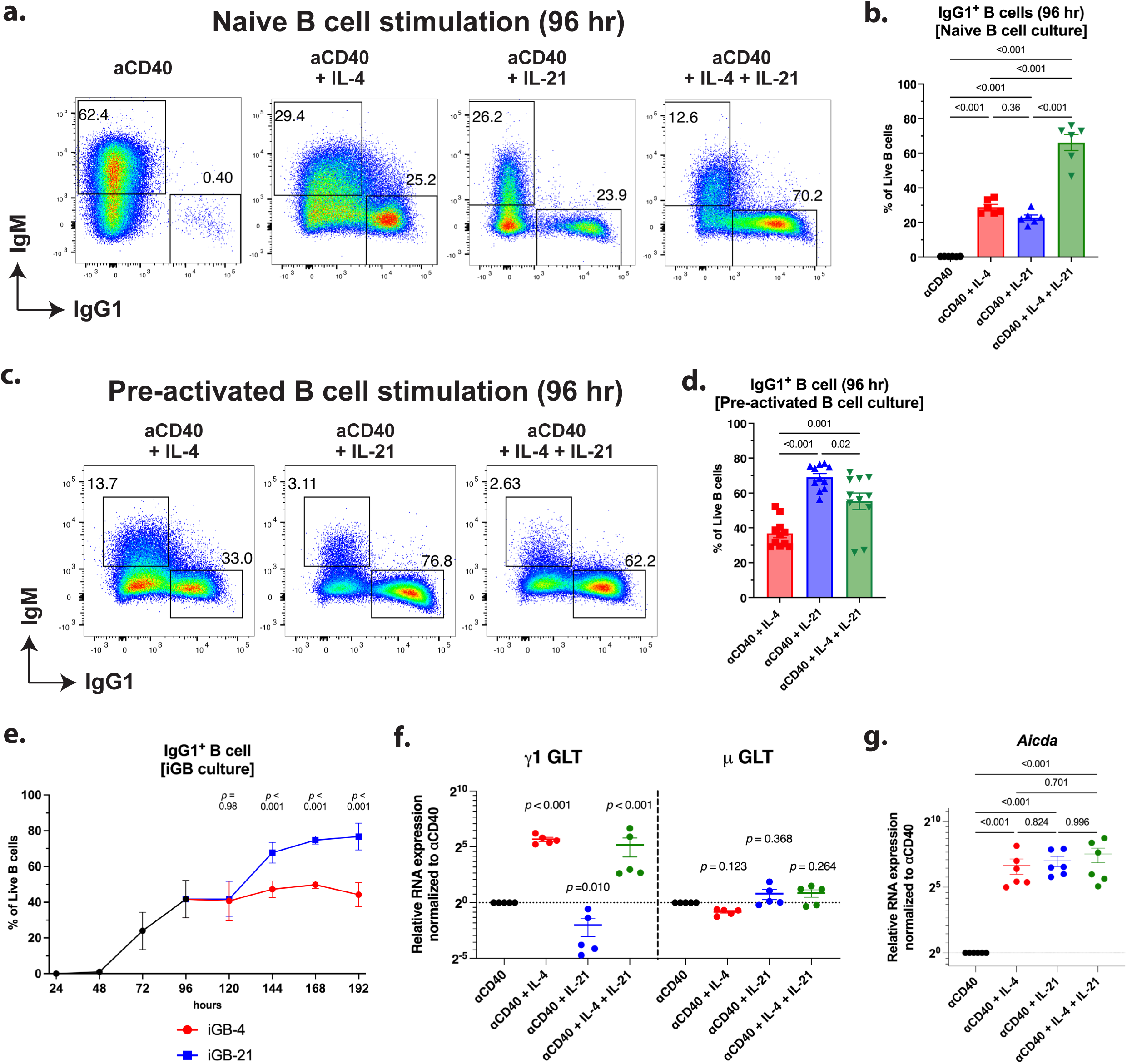
IL-21 induces expansion of IgG1^+^ cells *ex vivo* without driving *de novo* CSR. (a, c) Representative flow cytometry plot displaying IgM^+^ and IgG1^+^ B cell gating after 96 hours of *ex vivo* culture for (a) naïve B cell or (c) pre-activated B cells. Gating was adjusted for different stimulation conditions. (b, d) Percentage of IgG1-positive cells among live B cells after 96 hours of *ex vivo* culture for (b) naïve B cell or (d) pre-activated B cells. (e) Percentage of IgG1^+^ B cells among the live B cell population in the iGB culture. (f-g) Quantitative RT-PCR analysis for (f) germline transcript or (g) *Aicda* mRNA 48 hours after the culture of naïve primary B cells with the indicated combinations of stimulation. Each dot represents a biological replicate, and each biological replicate is the average of three technical replicates. The y-axis represents the expression of germline transcripts relative to that of B cells from the same animal stimulated with αCD40. Germline transcript expression for each sample was calculated by subtracting the (f) β-actin or (g) *Ubc* C_T_ (cycle threshold) value from the sample C_T_ value. For (b), (f), and (g), data were pooled from two independent experiments with *n* = 3 per experiment. For (d) and (e), data were pooled from three independent experiments with *n* = 3 or 4 per experiment. Data in (b), (d), (f) and (g) were analyzed with ANOVA in conjunction with Tukey’s multiple comparisons test. For (f), αCD40 was used as the control for statistical test. For (g), pairwise comparison for every condition was performed. Data in (e) were analyzed with Student’s unpaired t-test, with multiple comparisons correction applied using the Holm-Šídák method.

To investigate whether IL-21 directly influences CSR to IgG1, we evaluated sterile germline transcription (GLTs) from the Iγ1 promoter, which precede CSR to IgG1 and is often utilized as a proxy for the induction of CSR to IgG1^30^. While IL-4 increased the levels of γ1 GLT 30-fold above those observed in αCD40 alone, addition of IL-21 did not lead to a further increase in γ1 GLT (**Figure 3f**). RNA-seq data further indicated that IL-21, unlike IL-4, does not enhance transcription through the γ1 locus (**Supplementary Figure 2h**). As the expression of AID (encoded by *Aicda*) upon activation is a critical step in CSR^31,32^, we assessed transcription of *Aicda* mRNA by qPCR after 48 hours of culture. Addition of IL-4 and/or IL-21 on top of αCD40 enhanced *Aicda* expression compared to αCD40 alone. However, B cells stimulated with αCD40 + IL-4 or αCD40 + IL-21 showed similar levels of *Aicda* expression. Furthermore, adding IL-21 to αCD40 + IL-4 did not further elevate *Aicda* expression (**Figure 3g**). These observations suggest that IL-21 promotes expansion of the IgG1^+^ population without inducing γ1 GLT or enhancing AID expression beyond the levels induced by IL-4.

### IL-21 drives preferential proliferation of class-switched B cells *ex vivo*

The observation that IL-21 stimulation results in a pronounced expansion of the IgG1^+^ population without inducing γ1 GLT or AID expression led us to hypothesize that IL-21 expands IgG1^+^ B cell population by promoting the proliferation of pre-existing IgG1^+^ B cells rather than inducing *de novo* class switching to IgG1. Indeed, AID-deficient B cells, which cannot undergo class switch recombination and remain IgM^+^, proliferated significantly slower than AID-sufficient B cells in the iGB-21, but not in the iGB-4 culture (**Supplementary Figure 4a, b**).

To compare the proliferation of AID-sufficient and deficient B cells within the same microenvironment, we conducted a competitive *ex vivo* co-culture experiment. Naïve splenic B cells from AID-sufficient CD45.1^+^ mice and AID-sufficient or -deficient CD45.2^+^ mice were separately cultured on 40LB feeder cells with IL-4 for 96 hours. Subsequently, the CD45.1^+^ and CD45.2^+^ B cells were mixed in a 1:1 ratio and cultured in either the iGB-4 or the iGB-21 system (**Figure 4a**). In the iGB-4 culture, the relative proportion of CD45.1^+^ and CD45.2^+^ B cells remained largely unchanged regardless of AID genotype. However, in the iGB-21 culture, AID-sufficient CD45.1^+^ B cells far outcompeted AID-deficient CD45.2^+^ B cells (**Figure 4b, c**). Quantification of fold-change in cell number in iGB-21 culture revealed a markedly diminished ability of AID-deficient B cells to undergo proliferation relative to their WT counterparts (**Figure 4d**).

**Figure 4.**
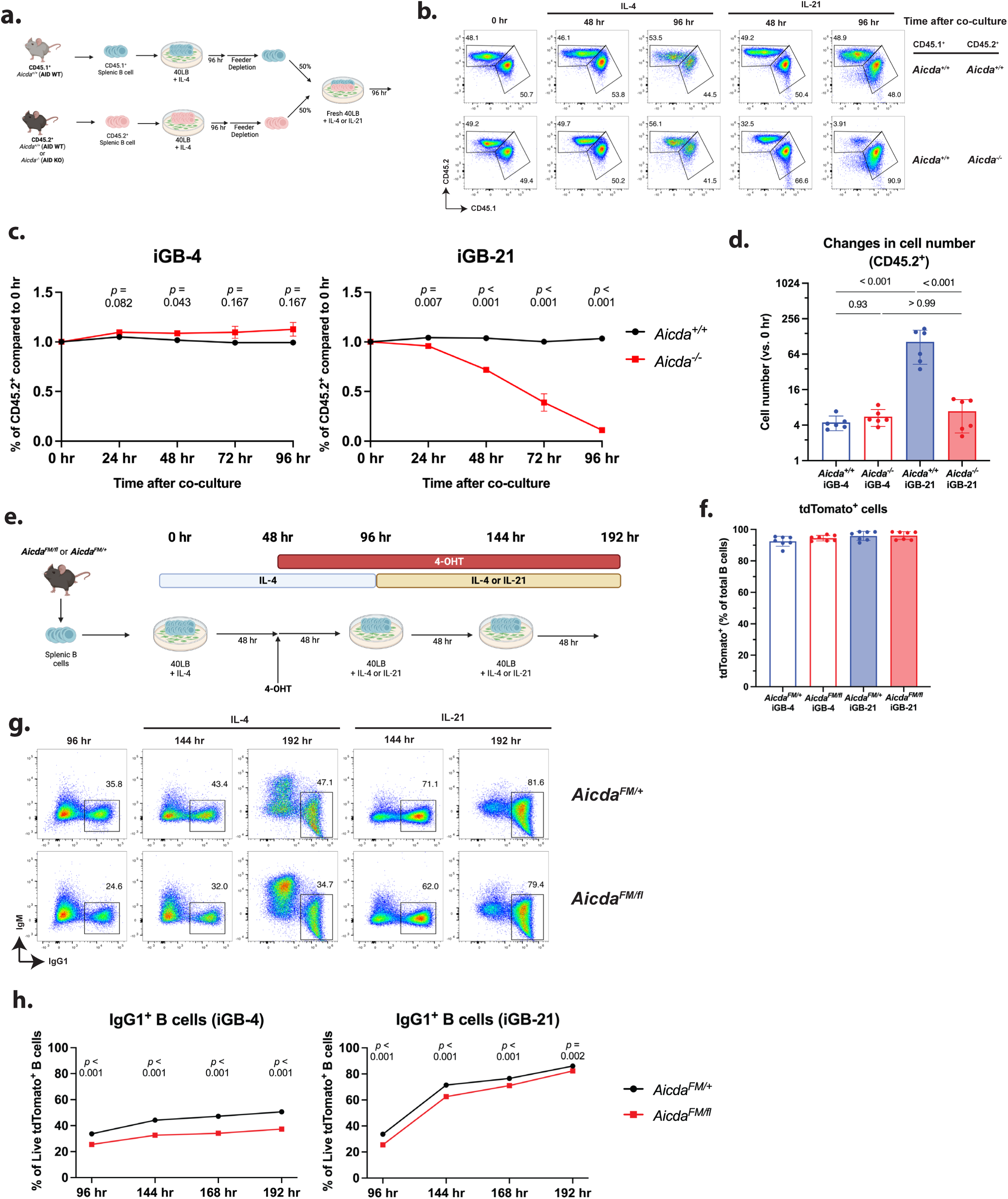
IL-21 drives proliferation of class-switched IgG1^+^ cells *ex vivo*. (a) Schematic representation of the competitive iGB culture setup. Naïve splenic B cells from CD45.1^+^ *Aicda^+/+^* mice and CD45.2^+^ *Aicda^+/+^* or *Aicda^-/-^* mice were initially cultured separately with the 40LB feeder cells and IL-4 (1 ng/mL) for 96 hours. Subsequently, feeder cells were depleted from the culture, and CD45.1^+^ and CD45.2^+^ B cells were mixed in equal proportions and cultured further with fresh 40LB feeder cells in the presence of IL-4 (1 ng/mL) or IL-21 (10 ng/mL). (b) Representative flow cytometry plot displaying the proportion of CD45.1^+^ and CD45.2^+^ cells in culture over time. (c) Changes in the percentage of the CD45.2^+^ population over time in the culture. The y-axis represents changes in the percentage of the CD45.2^+^ population relative to the start of the co-culture. (d) Changes in cell numbers of CD45.2^+^ B cells at 192 hours compared to the start of the co-culture (96 hours) under different stimulation conditions and genotypes. (e) Experimental setup illustrated schematically. Naïve splenic B cells from *Aicda^FM/fl^*or *Aicda^FM/+^* mice were initially cultured with the 40LB feeder cells and IL-4 (1 ng/mL). At 48 hours, 500 nM 4-OHT was added to the culture. At 96-hour, cells were cultured further with fresh 40LB feeder cells in the presence of IL-4 (1 ng/mL) or IL-21 (10 ng/mL). Feeder cells were replenished every 48 hours. (f) Percentage of tdTomato^+^ population among total B cells after 192 hours in culture. (g) Representative flow cytometry plot displaying the proportion of the IgG1^+^ population. (h) Changes in the percentage of IgG1^+^ population among tdTomato^+^ B cells over time in culture. Data were pooled from two independent experiments with *n* = 3 per experiment. Data in (c) and (h) were analyzed with Student’s unpaired t-test, with multiple comparisons correction applied using the Holm-Šídák method. Data in (d) were assessed with ANOVA in conjunction with Tukey’s multiple comparisons test.

To more precisely test the hypothesis that IL-21 leads to the preferential expansion of class-switched B cells rather than enhancing *de novo* class switching, we bred AID-flox mice^33^ with AID-fate mapping mice (*Aicda^creERT2/+^ Rosa26^Ai9/+^*) to generate the *Aicda^creERT2/fl^ Rosa26^Ai9/+^* mouse line, enabling tamoxifen-induced and AID-dependent deletion of the *Aicda* locus with concurrent tdTomato expression. For simplicity, *Aicda^creERT2/+^ Rosa26^Ai9/+^* mice will be referred to as *Aicda^FM/+^* and *Aicda^creERT2/fl^ Rosa26^Ai9/+^* mice as *Aicda^FM/fl^*. Naïve splenic B cells from *Aicda^FM/+^*or *Aicda^FM/fl^* mice were cultured on the 40LB feeder with IL-4, and 4-hydroxytamoxifen (4-OHT) was added at the 48-hour time point to stop further induction of CSR in *Aicda^FM/fl^* B cells. After 96 hours of culture, the cells were transferred to a fresh 40LB feeder layer and cultured using either iGB-4 or iGB-21 system (**Figure 4e**). Approximately 95% of *Aicda^FM/fl^*or *Aicda^FM/+^* B cells were tdTomato^+^ at the final timepoint in every experimental group (144 hours after 4-OHT treatment) (**Figure 4f**). Despite the loss of AID expression, 4-OHT treated *Aicda^FM/fl^* B cells underwent a significant expansion of IgG1^+^ population following IL-21 treatment. Strikingly, the difference in the frequency of IgG1^+^ B cells between *Aicda^FM/fl^*and *Aicda^FM/+^* B cells grew smaller over time in the presence of IL-21, but not with IL-4 treatment (**Figure 4g, h**). Collectively, these data imply that IL-21 preferentially expands class-switched IgG1^+^ B cells *ex vivo*.

Given that BCR cross-linking counteracts IL-21-dependent B cell apoptosis (**Figure 2a, b**), we hypothesized that different tonic BCR signaling strength could contribute to the observed IL-21 dependent proliferative advantage of IgG1^+^ B cells. Indeed, IgG1^+^ B cells exhibited higher levels of phosphorylated Syk and Erk than IgM^+^ B cells 72 hours after αCD40 + IL-4 stimulation, indicating stronger BCR signaling (**Supplementary Figure 5a, b**). As the BCR was not specifically engaged in this system, the difference is likely due to differences in tonic BCR signaling strength. To investigate the role of tonic BCR signaling in the selection of IgG1^+^ B cells, we used ibrutinib to inhibit Btk, a key signal transducer downstream of BCR (**Supplementary Figure 5c**). We observed that treatment with ibrutinib led to a significant reduction in the percentage of IgG1^+^ B cells in the iGB-21 culture, while it had no effect on the iGB-4 culture (**Supplementary Figure 5d**). Collectively, these results suggest that IL-21 synergizes with stronger BCR signaling in IgG1^+^ B cells to drive robust proliferation.

### B cell-specific deletion of *Il21r* leads to an impaired antibody response *in vivo*

Having found that IL-21 can differentially regulate B cell fates based on their activation status and BCR isotype, we sought to elucidate the implications and functional significance of these observations *in vivo*. We thus employed CRISPR/Cas9 technology to generate an *Il21r-flox* mouse strain to enable conditional deletion of *Il21r* (**Supplementary Figure 6a**). Retroviral transduction of Cre recombinase on splenic B cells from *Il21r-flox* mice led to a reduction in surface IL-21R expression to a level comparable to that in *Il21r*^-/-^ B cells (**Supplementary Figure 6b**). To specifically delete IL-21R in the mature B cells, we crossed *Il21r-flox* and *Cd23-cre* mice. B cells from the resulting *Il21r^fl/fl^::Cd23-cre* (*Il21r^ΔCd23^*) mice lacked IL-21-induced STAT1/3 phosphorylation, apoptosis, and expansion of IgG1^+^ B cells, confirming that deletion of exon 3 of *Il21r* generates a null allele (**Supplementary Figure 6c-g**).

To investigate the B cell-intrinsic role of IL-21 in the antibody responses, *Il21r^ΔCd23^* mice, along with control *Il21r^fl/fl^* mice (*Il21r^fl^*), were immunized with 4-hydroxy-3-nitrophenyl acetyl (NP)-conjugated chicken gamma globulin (NP-CGG) in alum, followed by longitudinal analysis of NP-specific IgM and IgG1 serum titers (**Figure 5a**). While the total serum IgM and IgG1 titer exhibited similar trends between the two groups over time (**Figure 5b, c**), *Il21r^ΔCd23^* mice displayed reduced levels of both high-affinity (anti-NP_(7)_) and total (anti-NP_(36)_) NP-specific IgG1 antibodies compared to *Il21r^fl^* control mice at all time points post-immunization (**Figure 5g, h)**. However, NP-specific IgM antibody levels in *Il21r^ΔCd23^*mice converged to control levels by day 21 (**Figure 5d, e**). The ratio of high-affinity to total-affinity antibody titer, indicative of affinity maturation, was notably diminished in *Il21r^ΔCd23^* mice for both IgM and IgG1 antibodies, with a more pronounced decrease observed for IgG1 (**Figure 5f, i**). Collectively, these data suggest that IL-21 plays a crucial role orchestrating optimal T-dependent antibody responses, with a notably greater impact on IgG1 compared to IgM responses, consistent with our *ex vivo* findings.

**Figure 5.**
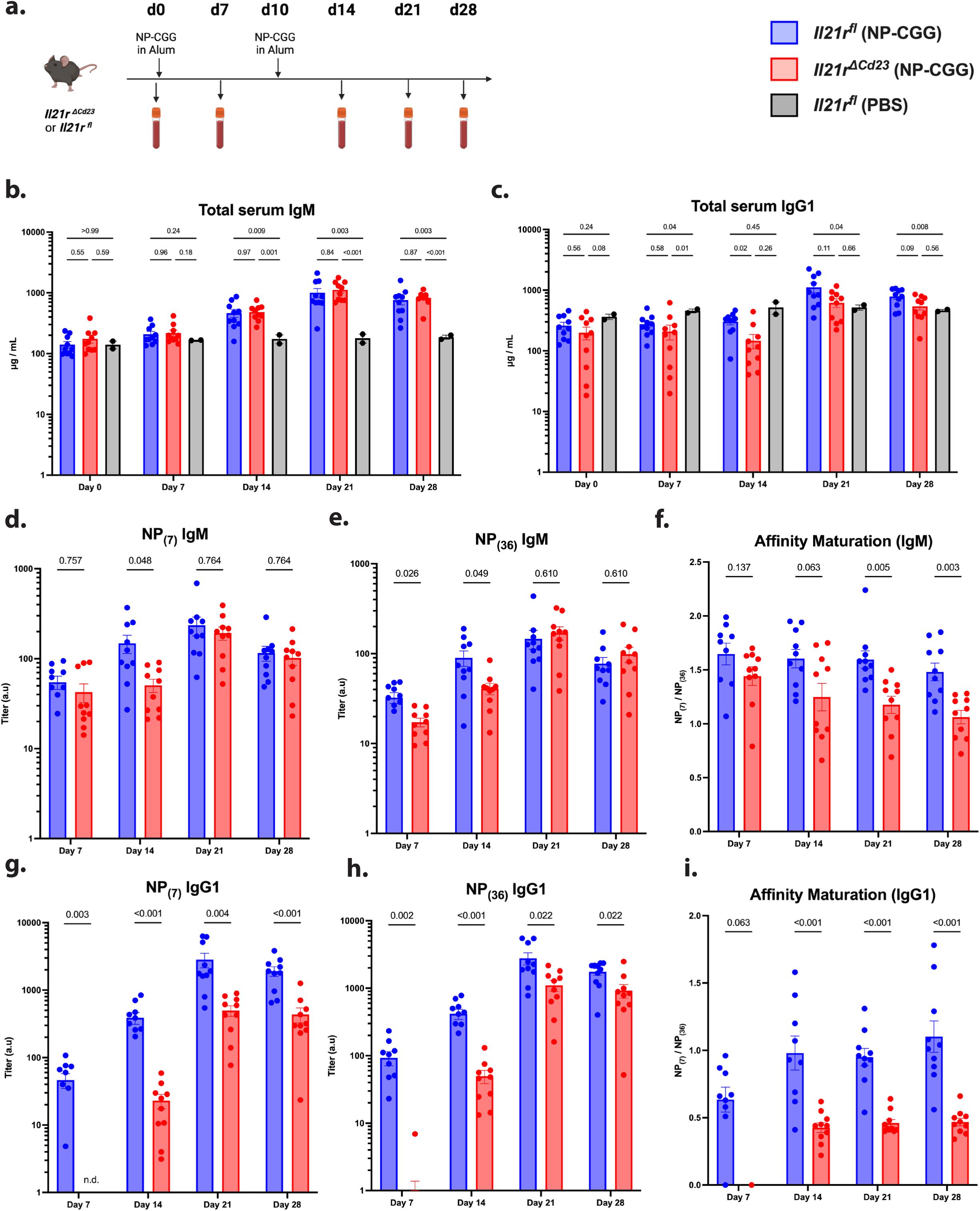
B cell-specific deletion of IL-21R impairs antigen-specific antibody responses. (a) Schematic representation of NP-CGG immunization and serum collection. *Il21r^ΔCd23^* mice and *Il21r^fl^* mice were immunized with 100 μg NP-CGG precipitated in Imject Alum or 100 μL PBS, followed by a booster immunization at day 10. Serum was collected from the mice every 7 days until day 28. (b, c) Concentration of total serum (b) IgM and (c) IgG1. The y-axis represents the antibody concentration in μg/mL. (d, e, g, h) Serum titer of (d) NP_(7)-_binding (high-affinity) IgM, (e) NP_(36)_-binding (total-affinity) IgM, (g) NP_(7)_-binding IgG1, and (h) NP_(36)_-binding IgG1. The y-axis represents the antibody titer in arbitrary units (a.u.). n.d. stands for not detected. (f, i) Ratio of NP_(7)_-binding to NP_(36)_-binding (f) IgM and (i) IgG1 titer. Different experimental groups are represented by distinct colors: *Il21r^fl^* mice immunized with NP-CGG (blue), *Il21r^ΔCd23^* mice immunized with NP-CGG (red), and *Il21r^fl^* mice injected with PBS (grey). Data were pooled from two independent experiments with *n* = 5 per experiment. Data in (b) and (c) were statistically analyzed with analyzed with two-way ANOVA in conjunction with Tukey’s multiple comparisons test. Data in (d), (e), (f), (g), (h), and (i) were statistically analyzed with Student’s unpaired t-test, with multiple comparisons correction applied using the Holm-Šídák method.

### IL-21 signaling in B cell is crucial for GC entry and maintenance

Given the pronounced defect in antibody affinity maturation in mice with B cell specific deletion of *Il21r*, we investigated the B cell intrinsic role of IL-21 signaling in GC responses. We immunized *Il21r^fl/fl^::Cd79a^creER/+^*(*Il21r^iΔMb1^*) mice with NP-conjugated keyhole limpet hemocyanin (NP-KLH) precipitated in alum adjuvant and cre activity was induced to delete the *Il21r* locus via oral gavage of tamoxifen dissolved in corn oil every two days, starting the day before immunization (**Supplementary Figure 7a**). We observed a significant reduction in the frequency and number of Fas^+^ GL7^+^ B cells in tamoxifen-treated *Il21r^iΔMb1^* mice compared to corn oil-treated control group (**Supplementary Figure 7b-d**).

To directly compare the GC responses of IL-21R-sufficient and -deficient B cells within the same animal, we generated chimeric mice whose B cells were derived from a 1:1 mixture of CD45.1^+^ *Il21r^fl^* and CD45.1/2^+^ *Il21r^ΔCd23^* bone marrow cells. We immunized the chimeric mice with NP-KLH precipitated in alum adjuvant (**Figure 6a)**. Remarkably, *Il21r^ΔCd23^* B cells exhibited a dramatic 10-fold reduction in percentages of Fas^+^ GL7^+^ B cells compared to *Il21r^fl^* B cells (**Figure 6b, c**), indicating that loss of IL-21 signaling in B cells severely compromises their ability to undergo a GC response.

**Figure 6.**
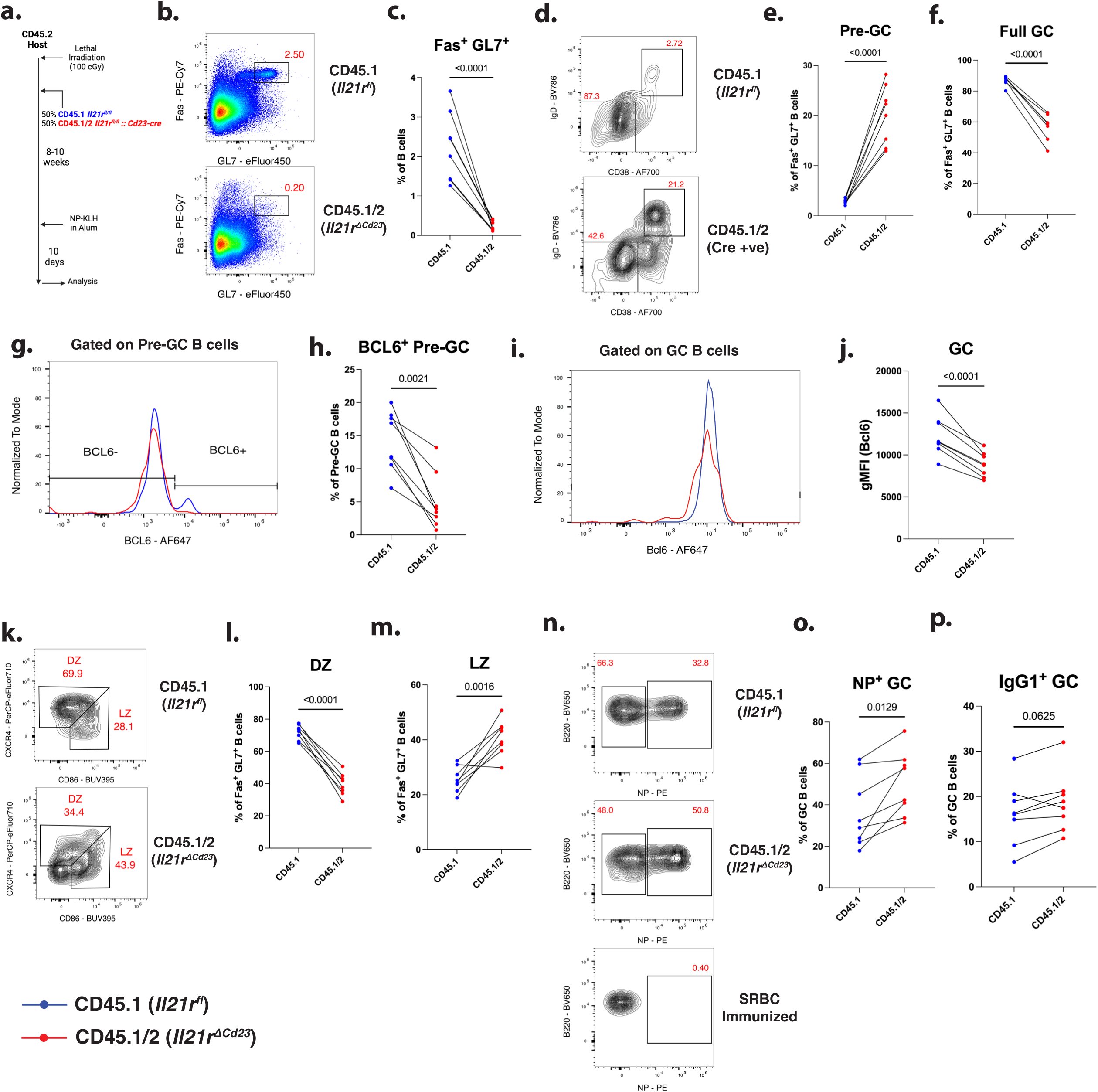
B cell-intrinsic IL-21 signaling is crucial for GC responses. (a) Schematic representation of NP-KLH immunization on mixed bone marrow chimeras reconstituted with *Il21r^fl^* (CD45.1^+^) and *Il21r^ΔCd23^*(CD45.1/2^+^) donor bone marrow cells. (b) Representative flow cytometry plot representing Fas^+^ GL7^+^ B cell gating. The plot is pre-gated on the live splenic B cell population. (c) Percentage of Fas^+^ GL7^+^ B cells among CD45.1^+^ or CD45.1/2^+^ splenic B cells. (d) Representative flow cytometry plot representing pre-GC (IgD^+^ CD38^hi^) and GC (IgD^-^ CD38^lo^) B cell gating. The plot is pre-gated on the live Fas^+^ GL7^+^ B cell population. (e, f) Percentage of (e) pre-GC and (f) GC B cells among CD45.1^+^ or CD45.1/2^+^ Fas^+^ GL7^+^ B cells. (g, i) Representative histogram of Bcl6 staining in the (g) pre-GC or (i) GC B cells populations. (h) Percentage of Bcl6^+^ cells within the pre-GC B cell population among CD45.1^+^ or CD45.1/2^+^ cells. (j) Geometric MFI of Bcl6 within the GC B cell population among CD45.1^+^ or CD45.1/2^+^ cells. (k) Representative flow cytometry plot representing DZ (CXCR4^hi^ CD86^lo^) and LZ (CXCR4^lo^ CD86^hi^) gating. (l, m) Percentage of (l) DZ and (m) LZ B cells within the live CD45.1^+^ or CD45.1/2^+^ Fas^+^ GL7^+^ B cell population. (n) Representative flow cytometry plot representing NP-specific GC B cells. The plot is pre-gated on the live GC B cell population. (o) Percentage of NP-specific B cells within the CD45.1^+^ or CD45.1/2^+^ GC B cell. (p) Percentage of IgG1^+^ B cells within the CD45.1^+^ or CD45.1/2^+^ GC B cell. Data were pooled from two independent experiments with *n* = 3 or 5 per experiment. Data in (c), (e), (f), (h), (j), (l), (m), (o), and (p) were statistically analyzed using ratio paired t-test.

While Fas^+^ GL7^+^ B cells have traditionally been considered as GC B cells, a recent study has revealed that this cell population consists of activated pre-GC B cells (Fas^int^ GL7^+^ CD38^+^ IgD^+^) and GC B cells (Fas^hi^ GL7^+^ CD38^-^ IgD^-^)^34^. Notably, while only 3% of *Il21r^fl^*-derived Fas^+^ GL7^+^ B cells exhibited a pre-GC B cell phenotype, this proportion increased to 20% in *Il21r^ΔCd23^*-derived Fas^+^ GL7^+^ B cells, implying impaired GC entry (**Figure 6d-f**). This trend was mirrored in tamoxifen-treated *Il21r^iΔMb1^* mice (**Supplementary Figure 7e-g**). Since transcription factor Bcl6 is required for GC entry^35^ and maintenance^36^, we examined Bcl6 expression in the pre-GC and GC B cell populations. Indeed, *Il21r^ΔCd23^* pre-GC B cells have a lower percentage of Bcl6^+^ cells compared to *Il21r^fl^* pre-GC B cells (**Figure 6g, h**). Moreover, *Il21r^ΔCd23^*GC B cells exhibited a substantial (approximately 30%) reduction in Bcl6 expression (geometric mean fluorescence intensity, gMFI) compared to their *Il21r^fl^*counterparts (**Figure 6i, j**). Taken together, these data suggest that loss of B cell-intrinsic IL-21 signaling impairs GC entry of B cells and diminishes the magnitude of GC responses, at least partly through regulating Bcl6 expression.

Quantitative alterations in the GC responses led us to further examine potential qualitative changes stemming from loss of IL-21 signaling. We observed that *Il21r^ΔCd23^*B cells exhibited a higher percentage of LZ cells and a lower percentage of DZ cells among Fas^+^ GL7^+^ B cells (**Figure 6k-m**). To investigate antigen-specific clones, we stained cells with NP-conjugated to phycoerythrin (PE). Surprisingly, *Il21r^ΔCd23^* B cells exhibited a notably higher percentage of NP^+^ B cells within the GC B cell population (**Figure 6n-o)**, possibly due to a more stringent selection for antigen specific clones in the absence of IL-21 signaling. This increased frequency of LZ cells and NP^+^ GC B cells was also observed in tamoxifen-treated *Il21r^iΔMb1^* mice (**Supplementary Figure 7h-l**). These findings suggest that IL-21 regulates GC dynamics by influencing the LZ/DZ distribution and the selection of antigen-specific clones.

### IL-21 promotes IgG1^+^ PB/PC cell selection, but not IgG1^+^ GC B cells *in vivo*

CSR is believed to occur predominantly in the pre-GC state, yet GCs are dominated by class-switched clones over time, suggesting the existence of mechanisms fostering the expansion of these clones^2,37^. Given our finding that IL-21 selectively promotes the outgrowth of IgG1^+^ B cells *ex vivo* (**Figure 4**), we posited IL-21 as a driver for this expansion. Contrary to our expectation, *Il21r^ΔCd23^* and *Il21r^fl^* B cells exhibited comparable percentage of IgG1^+^ cells within the GC (**Figure 6p**). Given the putative role of IL-21 in PB and PC differentiation^22^, we next examined the splenic PB/PC (TACI^+^ CD138^+^) population in chimeric mice. Indeed, we observed a significantly lower frequency of *Il21r^ΔCd23^*-derived PB/PC compared to *Il21r^ΔCd23^*-derived splenic B cells, indicating that IL-21 promotes PB/PC differentiation (**Figure 7a-b**). Additionally, among *Il21r^ΔCd23^*-derived PB/PCs, there was a higher percentage of IgM^+^ cells and a lower percentage of IgG1^+^ cells compared to their *Il21r^fl^*counterparts (**Figure 7c-f**). Furthermore, *Il21r^ΔCd23^*-derived PB/PC exhibited a lower percentage of NP-specific cells (**Figure 7g, h**), and among these, a lower percentage expressed IgG1, and a higher percentage expressed IgM (**Figure 7i, j**). These findings suggest that IL-21 promotes not only PB/PC differentiation but also the positive selection of IgG1^+^ PB/PC, while not affecting IgG1^+^ GC B cells, *in vivo*.

**Figure 7.**
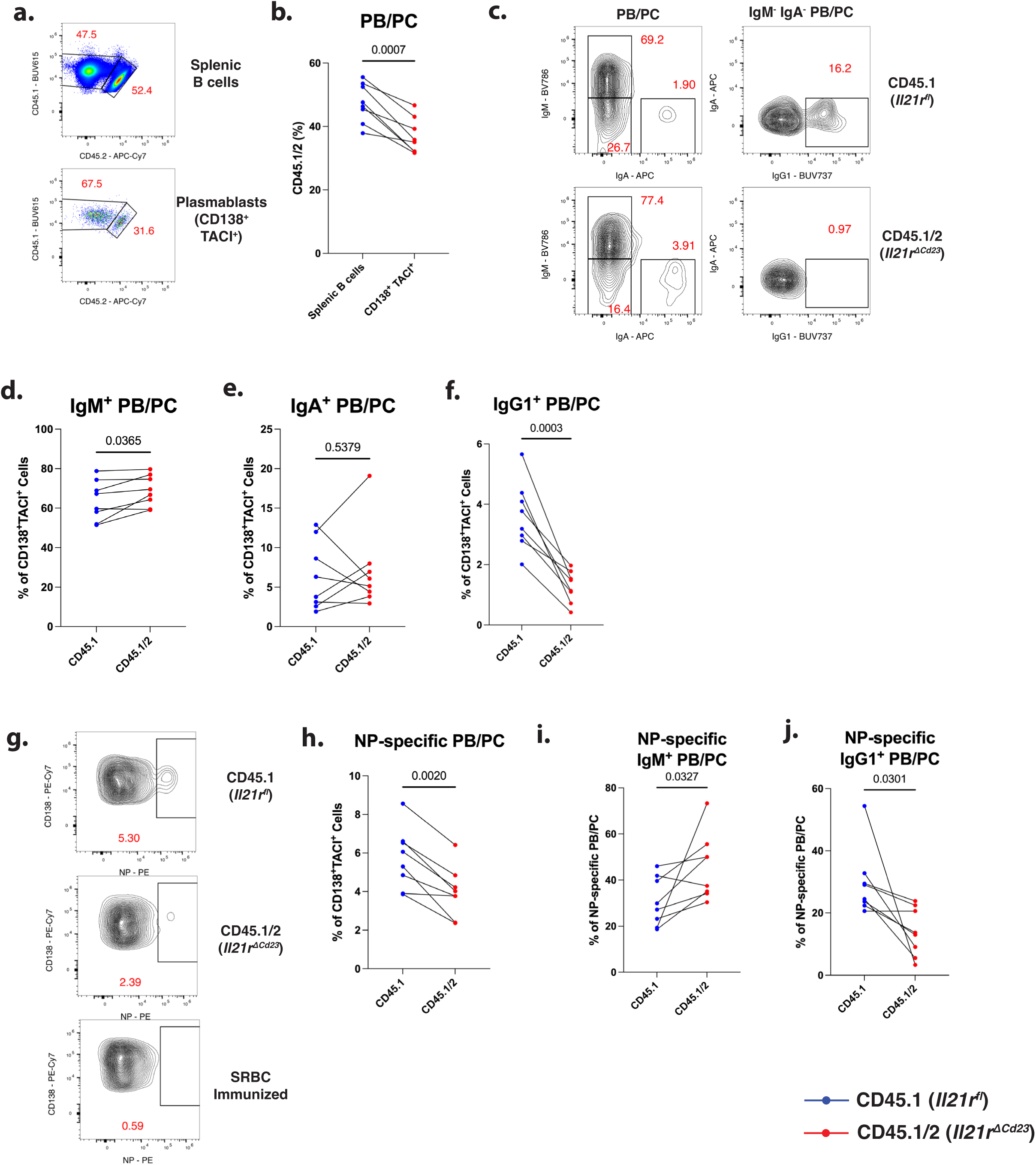
IL-21 promotes the selection of IgG1^+^ plasmablasts/plasma cells (PB/PCs). (a) Representative flow cytometry plot representing the gating of CD45.1^+^ or CD45.1/2^+^ cells within the splenic B cell (upper) or PB/PC (CD138^+^ TACI^+^ live singlets) (lower) populations. (b) Percentage of CD45.1/2^+^ donor-derived cells within the splenic B cell or PB/PC populations. (c) Representative flow cytometry plot representing IgM^+^, IgA^+^, and IgG1^+^ PB/PC gating. (d-f) Percentage of (d) IgM^+^, (e) IgA^+^, or (f) IgG1^+^ cells among CD45.1^+^ or CD45.1/2^+^ PB/PCs. (g) Representative flow cytometry plot representing NP-specific PB/PCs. (h) Percentage of NP-specific cells within the CD45.1^+^ or CD45.1/2^+^ PB/PCs. (i, j) Percentage of (i) IgM^+^ or (j) IgG1^+^ cells within the CD45.1^+^ or CD45.1/2^+^ NP-specific PB/PCs. Data were pooled from two independent experiments with *n* = 3 or 5 per experiment. Data in (b), (d), (e), (f), (h), (i), and (j) were statistically analyzed using ratio paired t-test.

## DISCUSSION

Since the discovery of IL-21 and its receptor around 25 years ago^15,38^, a huge body of literature has shed light on their crucial roles in supporting GC responses and how dysregulation of IL-21 signaling influences the pathogenesis of many autoimmune diseases^13^. Despite the breadth of past research, several aspects of IL-21 biology remain unresolved. These include the precise differentiation step at which IL-21 signaling is critical and the extent to which the IL-21-dependent effects are attributable to a B cell-intrinsic requirement of this cytokine. Results presented above provide unequivocal evidence for IL-21 signaling being a critical component of the GC response, demonstrate the existence of the pre-GC as a transition state for B cells prior to commitment to a full-fledged GC reaction, and illustrate a multifaceted role of IL-21 in promoting in either cell death or proliferation in a context-dependent manner.

We have found that naïve splenic B cells exposed to IL-21 in culture upregulate expression of the pro-apoptotic gene *Bcl2l11* and undergo apoptosis. Notably, strong BCR signaling confers protection against IL-21-induced apoptosis. We propose that this *ex vivo* phenomenon recapitulates an *in vivo* checkpoint that ensures elimination of bystander non-specific B cells in the vicinity of T_FH_ cells in lymphoid follicles. This control mechanism is crucial because IL-21 is capable of exerting effects beyond immunological synapses in a paracrine manner^39^. Additionally, recent research indicates that GC B cells modulate IL-21 binding through decreased heparan sulfate sulfation^40^, highlighting the critical importance of tightly regulating IL-21 activity within the GCs. Whether perturbation of such a step promotes the generation of autoreactive antibodies and increases the propensity for autoimmune syndromes awaits additional studies.

In contrast to the IL-21-induced apoptosis in naïve B cells, activated B cells have a markedly different response to the cytokine. Pre-stimulation of B cells with either αCD40 + IL-4 or growth of splenic B cells on 40LB feeder cells supplemented with IL-4 primes them for robust IL-21-dependent proliferation. This expansion of activated B cells by IL-21 appears to be dependent on the isotype of BCR expressed on B cells. IgG1^+^ B cells are significantly more responsive to IL-21 than IgM^+^ cells, as clearly demonstrated by a failure of AID-deficient B cells in effectively competing with wild-type B cells in *ex vivo* cultures. The preferential expansion of IgG1^+^ B cells might be due to a stronger tonic signaling from their BCR compared to IgM^+^ B cells. This is supported by the fact that inhibiting BCR signaling significantly mutes IL-21-dependent proliferation of IgG1^+^ B cells. This finding is consistent with previous reports highlighting the synergistic interplay among IL-21R, BCR, and CD40 signaling pathways in B cell activation^39–41^.

The *Il21r-flox* mouse model generated in this study allowed us to unequivocally assess the B cell-intrinsic requirement of IL-21 signaling in GC dynamics *in vivo*. One of the most prominent observations from the mouse model is an impaired transition from the pre-GC to the GC state in the absence of IL-21 signaling in B cells. It is becoming evident that upon BCR engagement in the follicles, B cells transit through a pre-GC checkpoint^42^ wherein the fitness and appropriate antigen-engagement of a B cell is evaluated prior to entry into the energetically demanding^43^ and generally genome-destabilizing environment of the GCs^44^. We suggest that IL-21 is one of the effectors of this pre-GC to GC checkpoint. One possible mechanism by which IL-21 influences this transition is through the upregulation of Bcl6^19^. It is important to note that B cells upregulate IL-21R expression upon the exposure to IL-21, suggesting that a positive feedback loop might exist and could be crucial for B cells to achieve the level of Bcl6 necessary to pass the checkpoint.

Loss of IL-21 signaling led to reduced Bcl6 expression not only in pre-GC B cells but also in GC B cells, suggesting premature exit from GCs or a failure to establish a fully mature GC. This could explain the observed reductions in antigen-specific serum antibody titers and compromised affinity maturation following immunization. The *Il21r*-deficient GC B cells have intriguing differences from *Il21r*-sufficient GC B cells. Firstly, *Il21r*-deficient GC B cells predominantly occupy the LZ while wild type B cells predominantly populate the DZ. This altered GC dynamics, also noted in previous studies^45–48^, probably reflects a lack of appropriate T-cell help in the GCs and/or a failure to sustain DZ proliferation. Second, *Il21r*-deficient GCs have a heightened proportion of antigen-specific B cell clones relative to control mice. This finding, resonating with a previous study on B cell specific STAT3 deficiency^49^, has two non-mutually exclusive possible explanations. First, loss of IL-21 signaling may enforce a more stringent selection for antigen specificity. Given that IL-21 is implicated in the positive selection of B cells engaged with T_FH_ cells^40,41,46^, it is plausible that the affinity of BCR emerges as a potent driving force for positive selection in the absence of IL-21 signaling. This hypothesis gains support from recent research indicating that exogenous IL-21 introduction during GC responses reduces the proportion of high-affinity antigen-specific GC B cells, suggesting a less stringent positive selection in the presence of excessive IL-21^40^. Second, it is possible that lack of IL-21 signaling blocks differentiation of GC B cells into PCs, leading to accumulation of high-affinity clones, which are known to be preferentially selected into the PC pool. Indeed, increased proportion of antigen-specific GC B cells in *Il21r*-deficient GCs is accompanied by a reduction in the proportion of antigen-specific PB/PC.

There is an apparent disconnect between the IL-21-dependent expansion of class-switched B cells *ex vivo* and the relatively normal frequency of IgG1^+^ *Il21r*-deficient B cells within GCs relative to controls *in vivo*. However, we found a substantial reduction in the percentage of IgG1^+^ cells among PB/PCs in *Il21r*-deficient B cells *in vivo*. It is generally believed that high-affinity GC B cells are preferentially selected into the PC lineage through both BCR signaling and T_FH_ cell-derived signals^50,51^. Our observations suggest that stronger tonic BCR signaling in IgG1^+^ cells synergize with T_FH_-derived IL-21 to preferentially select IgG1^+^ cells into the PB/PC pool. This could provide an additional explanation for the predisposition of IgG1^+^ memory B cells to differentiate into PCs^52^.

In summary, this study provides additional evidence supporting the critical role of IL-21 signaling in B cells in orchestrating GC responses and facilitating affinity maturation. Our findings further suggest that IL-21 may influence B cell fate decisions based on their activation status. The novel *Il21r-flox* mouse model developed herein offers a valuable tool to further decipher the involvement of IL-21 signaling not only in B cells but also in T cells during GC responses. Furthermore, given the implication of IL-21 in various autoimmune diseases such as systemic lupus erythematosus, rheumatoid arthritis, and type 1 diabetes^53^, this model holds promise as a resource for uncovering their underlying pathobiology and identifying potential therapeutic targets.

## MATERIAL AND METHODS

### Mice

*Il21r-flox* mice were generated utilizing CRISPR/Cas9 genome editing technology at the MSKCC transgenic core facility. Guide RNAs targeting upstream and downstream of the exon 3 of *Il21r* locus and donor oligonucleotides containing *loxP* sites were used to generate the mouse line. Wild type C57BL6/J mice were procured from the Jackson Laboratory (JAX# 000664). Additional mouse strains used in this study were generously provided as follows: *Aicda^-/-^* (AID knock-out) mice from Dr. T. Honjo, *Aicda^cre/cre^*(AID-cre) mice from Dr. M. Nussenzweig (JAX #007770), *Aicda^fl/fl^*(AID-flox) mice from Dr. D. Bhattacharya (JAX #035763), *Aicda^creERT2/creERT2^*(AID-creERT2) mice from Dr. C.-A. Reynaud (JAX #033897), *Rosa26^LSL-tdTomato/LSL-tdTomato^* (*Rosa26^Ai9/Ai9^*) mice from Dr. H. Zeng (JAX #007909), *Cd79a^creERT2/creERT2^* (Mb1-creERT2) mice from Dr. E. Hobeika (JAX #033026), Cd23-cre mice from Dr. M. Busslinger (JAX #028197), *Stat3^fl/fl^* (Stat3-flox) mice from Dr. X.-Y. Fu (JAX #016923), *Bcl2l11^-/-^* (Bim KO) mice from Dr. S. J. Korsmeyer, and C57BL/6-CD45.1^STEM^ (CD45.1) mice from Dr. D. T. Scadden. All mice were maintained in specific-pathogen free conditions and handled in compliance with the guidelines stipulated by MSKCC Research Animal Resource Center (RARC) and Institutional Animal Care and Use Committee (IACUC).

### Naïve B cell purification and *ex vivo* primary B cell culture

Mouse spleens were dissociated through a 70μm cell strainer and single cell suspension was prepared in B cell media (RPMI-1640, 15% FBS, 1% Pen-Strep, 1% L-glutamine, and 55 mM β-mercaptoethanol). Cells were collected by centrifugation and resuspended in red blood cell lysis buffer (0.15 M NH_4_Cl, 0.01 M KHCO_3_, and 100 μM EDTA in 1X PBS without Ca^2+^ and Mg^2+^). This mixture was incubated at room temperature for 5 minutes to achieve erythrocyte lysis and then neutralized with B cell media. After another round of centrifugation, the cell pellets were resuspended in 5 mL of DNase I reaction buffer (60 μg/mL DNase I (Roche) and 0.1% BSA in 1X PBS with Ca^2+^ and Mg^2+^). The suspension was incubated at room temperature for 20 minutes with gentle shaking to facilitate the digestion of extracellular DNA. Following enumeration using a hemocytometer, naïve B cells were purified by negative selection using anti-CD43 magnetic beads (Miltenyi) according to manufacturer’s protocol. Naïve splenic B cells were cultured in B cell media at a density of 1 × 10^6^ cells/mL. B cells were stimulated under various conditions as listed below, unless otherwise specified: αCD40 mAb (Invitrogen or BioLegend, Clone HM40-3, 1 μg/mL), IL-4 (R&D Systems, 12.5 ng/mL), IL-21 (R&D Systems, 50 ng/mL), αIgM mAb (F(ab’)2-Goat anti-Mouse IgM, Invitrogen, 1 μg/mL).

### Hydrodynamic gene transfer

The cDNA encoding murine IL-21 was cloned into the pCMV3 expression vector (Sino Biological), and 20 μg of plasmid in 2 mL of PBS was injected into the tail vein within 6 to 8 seconds. Mice that received injections of pCMV3-IL-21 or the empty pCMV3 vector were analyzed 7 days post-injection.

### Retroviral construction and transduction of primary B cells

For retrovirus production, HEK293T cells were used. HEK293T cells were cultured in 293T media (DMEM, 10% FBS, 1% Pen-Strep, 1% L-Glutamine) in 10-cm tissue culture dishes and split every three days 1:15 into new 10-cm tissue culture dishes. To transfect a single 10-cm dish culture, a transfection mix was prepared by mixing 1.35 mL of Opti-MEM (ThermoFisher) with 150 μL of PEI (1 mg/mL) and this mixture was added dropwise to 1.5 mL of Opti-MEM containing 20 μg of pCL-Eco, and 30 μg of the retroviral plasmid. After a brief vortexing and incubating the 3 mL transfection mix for 20 minutes at room temperature to form DNA-PEI complexes, the cell culture media from confluent HEK293T cells were aspirated, and the transfection mix was carefully added to the cells, followed by incubation at 37°C for 4 hours. Subsequently, the transfection mix was aspirated, and replaced with fresh 10 mL of 293T media. At 48- and 72-hours post-transfection, supernatants were collected and combined. The viral particles were concentrated using Lenti-X Concentrator (Takeda) as per the manufacturer’s instructions and stored at -80°C in aliquots.

For retroviral transduction, naïve splenic B cells were initially cultured in the presence of αCD40 and IL-4, then spinfected twice, at 24 hours and 48 hours of culture. During this process, concentrated viral supernatant was mixed with polybrene (reaching a final concentration of 8 μg/mL) and added to the B cell culture. The culture was then subjected to centrifugation at 800 x g and 32°C for 90 minutes. Subsequently, half of the culture’s supernatant was removed, and an equal volume of fresh B cell media and was added along with αCD40 and IL-4. Cells were analyzed 48 hours after the second retroviral spinfection.

### iGB culture

40LB cells are generous gift of Dr. D. Kitamura. 40LB cells were cultured in 40LB media (DMEM, 10% FBS, 2 mM Sodium Pyruvate, 1% NEAA, 1% Pen-Strep, 1% L-Glutamine) in 10-cm tissue culture dishes and split every three days to maintain a density of 5 × 10^5^ cells per 10-cm tissue culture dish. For the iGB culture, confluent 40LB cells were harvested and irradiated at 100 Gy using a cesium irradiator (J L Shepherd & Associates), after which they were plated on 6-well tissue culture plates at a density of 5 × 10^5^ cells in 2 mL of 40LB media per well. Following overnight incubation at 37°C, the culture media was aspirated, and the adherent irradiated 40LB cells were co-cultured with 5 × 10^4^ naïve splenic B cells in 6 mL of B cell media containing IL-4 (R&D Systems, 1 ng/mL) for 96 hours. Cells were harvested by collecting the culture supernatant and incubating the wells with 3 mL of MACS buffer until cell detachment. The detached cells were pooled with the harvested supernatant, and each well was washed once more with 3 mL of MACS buffer. For secondary cultures, 1 × 10^5^ B cells were added to each well in 6 mL of B cell media containing either IL-4 (R&D Systems, 1 ng/mL) or IL-21 (R&D Systems, 10 ng/mL) with fresh irradiated 40LB feeder cells. Feeder cells were replenished every 48 hours, and when needed, 4-OHT (Sigma, 500 nM) or ibrutinib (Selleckchem, 100 nM) was introduced into the culture. For feeder cell depletion, harvested cells were incubated with anti-H-2Kd antibody (2.9 μg/mL in MACS buffer), which binds the feeder cells, at room temperature for 20 minutes. After washing, the pellet was resuspended with streptavidin particle cocktail (100 μL MACS buffer + 50 μL BD IMag Streptavidin Particles Plus-DM (BD) per 10^7^ cells), incubated at room temperature for 20 minutes. After the incubation, the labeled cells were placed on the magnetic stand and the unlabeled cells were harvested.

### Flow cytometry sample preparation

Mouse splenocytes were harvested as described earlier for naïve B cell purification. For general flow cytometry analysis, cells were washed with 1X PBS with Ca^2+^ and Mg^2+^ and stained with Zombie Red, Zombie NIR, or Zombie Yellow fixable viability dye (BioLegend) 1:1000 in 1X PBS with Ca^2+^ and Mg^2+^ for 15 minutes at room temperature in the dark. Subsequently, cells were washed with FACS buffer (2.5% FBS in 1X PBS with Ca^2+^ and Mg^2+^) and stained with anti-CD16/32 monoclonal antibody (BD or Cytek, Clone 2.4G2, 1:50 in FACS buffer) for 20 minutes at 4°C in the dark to block Fc receptors. After a wash with FACS buffer, cells were stained with the surface antibody cocktail in FACS buffer for 20 minutes at 4°C in the dark. The cells were washed twice with FACS buffer, resuspended in FACS buffer, and analyzed.

For intracellular staining of active caspase-3, prior to incubation with surface-staining antibodies, cells were fixed with BD CytoFix/CytoPerm Fixation Buffer for 20 minutes at 4°C in the dark. Following two washes with 1X BD Perm/Wash buffer, cells were stained with AlexaFluor 647-rabbit anti-active caspase-3 antibody (BD, 1:20 in 1X BD Perm/Wash buffer) for 30 minutes at 4°C in the dark. Cells were washed twice with 1X BD Perm/Wash buffer and analyzed after resuspension in FACS buffer.

For intracellular staining of phosphorylated proteins or Bim, cells were fixed with an equivolume of 4% PFA (pre-warmed at 37°C, final concentration 2% PFA) at 37°C for 30 minutes. Fifteen minutes before the end of incubation, Zombie Red fixable viability dye diluted in B cell media with 2% PFA was added to a final concentration of 1:1,000. After washing twice with FACS buffer, cells were stained with anti-CD16/32 monoclonal antibody followed by the surface staining cocktail, as described in the general sample preparation. After washing twice with FACS buffer, cells were permeabilized with Perm Buffer III (BD, pre-chilled at -20°C) at 4°C for 30 minutes. Following another wash with FACS buffer, cells were stained with the anti-phosphoprotein or Bim antibody cocktail (1:50 in FACS buffer) overnight at 4°C in the dark. Cells were washed twice with FACS buffer, resuspended in FACS buffer, and analyzed. All samples were analyzed using either BD LSR II or Cytek Aurora flow cytometers and analyzed with FlowJo (v. 10.9.0).

### Quantitative RT-PCR (qRT-PCR)

For analysis of germline transcript or *Aicda* mRNA expression, purified naïve splenic B cells were stimulated, and total RNA was extracted using the Zymo RNA miniprep kit (ZymoResearch) following manufacturer’s protocol. The concentration of purified RNA was determined using a Nanodrop (ThermoFisher). Subsequently, 1 μg of total RNA from each sample was used for reverse transcription. The reverse transcription reaction was carried out using the SuperScript IV reverse transcriptase kit (ThermoFisher) with random hexamers as primers, following manufacturer’s protocol.

For quantitative PCR (qPCR) of germline transcripts, each reaction had a total volume of 10 μL and contained the following components: 5 μL of 2X PowerUP SYBR Green master mix (ThermoFisher), 2 μL of the reverse transcribed product, 0.5 μL of a 10 μM forward primer (final concentration 500 nM), 0.5 μL of a 10 μM reverse primer (final concentration 500 nM), and 2 μL of DEPC-treated water. For the background control, 2 μL of DEPC-treated water was used instead of the reverse transcribed product. For μ-GLT, the forward primer sequence used was 5’-CTC TGG CCC TGC TTA TTG TTG-3’, the reverse primer sequence was 5’-GAA GAC ATT TGG GAA GGA CTG ACT-3’; Tm for both is 60°C. For γ1-GLT, the forward primer sequence was 5’-GGC CCT TCC AGA TCT TTG AG-3’, the reverse primer sequence was 5’-GGA TCC AGA GTT CCA GGT CAC T-3’; Tm for both is 58°C. As a control, β-actin was used, with the forward primer sequence 5’-TGC GTG ACA TCA AAG AGA AG-3’, the reverse primer sequence 5’-CGG ATG TCA ACG TCA CAC TT-3’, Tm is 55°C.

For qPCR of *Aicda* mRNA, each reaction had a total volume of 10 μL and contained the following components: 5 μL of 2X TaqMan Fast Advanced Master Mix (Applied Biosystems), 1 μL of the reverse transcribed product, 0.5 μL of TaqMan Gene Expression Assay Probe (20X) [Mm01184115_m1 for *Aicda*, *Mm01201237_m1* for the Ubc control], and 3.5 μL of DEPC-treated water.

Each qPCR reaction was performed in technical triplicates, and the reactions were set up in an optical 384-well plate. The PCR was run on a QuantStudioTM 6 Real-Time PCR instrument (ThermoFisher). The PCR cycling conditions for germline transcript analysis were as follows: UDG activation at 50°C for 2 minutes, polymerase activation at 95°C for 2 minutes, followed by 40 cycles of denaturation at 95°C for 15 seconds, annealing at the primer-specific Tm for 15 seconds, and elongation at 72°C for 1 minute. A dissociation curve analysis was carried out with the following steps: 95°C for 15 seconds (ramp rate = 1.6°C/second), 60°C for 1 minute (ramp rate = 1.6°C/second), and 95°C for 15 seconds (ramp rate = 0.15°C/second). The PCR cycling conditions for *Aicda* expression analysis were as follows: UDG activation at 50°C for 2 minutes, polymerase activation at 95°C for 2 minutes, followed by 40 cycles of denaturation at 95°C for 3 seconds and annealing/extending at 60°C for 30 seconds. The cycle threshold (C_T_) values were calculated using QuantStudio Software (ThermoFisher), and the relative expression of germline transcripts was determined using the 2^-ΔΔCT^ method with β-actin or *Ubc* as the internal control.

### RNA-sequencing and analysis

For RNA-sequencing experiments, total RNA was extracted using either the Zymo RNA miniprep kit (ZymoResearch) or the PicoPure™ RNA Isolation Kit (Applied Biosystems), following the manufacturer’s protocols. To ensure the quality of the RNA preparations, an assessment was performed using Tapestation RNA Screentape (Agilent). For the RNA samples that met the quality criteria, either mRNA enrichment or rRNA depletion was carried out using the NEBNext® Poly(A) mRNA Magnetic Isolation Module (New England Biolabs) or RiboCop rRNA Depletion Kits (Lexogen). Subsequent library preparation was conducted using the xGen™ Broad-Range RNA Library Preparation Kit (Integrated DNA Technologies), in conjunction with xGen UDI Primers (Integrated DNA Technologies). The quality of the prepared libraries was assessed using Tapestation D1000 Screentape (Agilent). Following library quality control, the libraries that met the necessary standards were sequenced. Sequencing was carried out by the MSKCC Integrative Genomics Operation (IGO) using the NovaSeq 6000 system (Illumina). Each library was sequenced at 30-40 million PE100 reads. RNA-sequencing reads were quantified using Salmon^54^ (v1.9.0). The mouse GRCm38 (mm10) reference transcriptome served as the basis for indexing Subsequently, the Salmon quantification files were processed using the R package tximport^55^ (v1.28.0). Normalization and differential gene expression analysis were conducted employing the R package DESeq2^56^ (v1.40.2). Furthermore, gene set enrichment analysis focusing on KEGG pathways was performed utilizing the R package fgsea^57^ (v1.26.0).

### Mixed Bone Marrow Chimera

Mixed bone marrow chimeras were generated by lethal irradiation (1000 cGy) of host CD45.2 mice and subsequent reconstitution with a 1:1 mix of bone marrow cells derived from *Il21r^fl/fl^* (CD45.1) and *Il21r^fl/fl^*::*Cd23-cre* (CD45.1/2) hosts. The mice were analyzed 8–12 weeks following the bone marrow transplant.

### Immunization

For SRBC immunization, SRBCs (Innovative Research) were washed with sterile 1X PBS with Ca^2+^ and Mg^2+^ and enumerated using a hemocytometer. SRBCs were then diluted to a concentration of 5 × 10^9^ cells/mL using sterile 1X PBS with Ca^2+^ and Mg^2+^. Each mouse received an intraperitoneal injection of 100 μL of the diluted SRBCs, equivalent to 5 × 10^8^ cells. For booster immunization, an equal amount of diluted SRBCs was intraperitoneally injected 10 days after the initial immunization.

For immunization with NP-conjugated carrier antigen, lyophilized NP-CGG or NP-KLH (Biosearch Technologies) was reconstituted with sterile 1X PBS with Ca^2+^ and Mg^2+^ to a concentration of 3 mg/mL. To prepare the immunization mixture, the reconstituted NP-CGG or NP-KLH was transferred to a small beaker, and twice the volume of Imject Alum (ThermoFisher) was added dropwise, resulting in a final NP-CGG or NP-KLH concentration of 1 mg/mL. NP-CGG or NP-KLH was emulsified in alum by magnetic stirring at room temperature for 30 minutes. For NP-CGG or NP-KLH immunization, each mouse was intraperitoneally injected with 100 μL of the NP-CGG or NP-KLH emulsion in alum, equivalent to 100 μg of NP-CGG or NP-KLH. Booster immunizations followed the same procedure, with an equal amount of NP-CGG or NP-KLH emulsion in alum being intraperitoneally injected 10 days after the initial immunization.

### ELISA

ELISA was conducted employing MaxiSorp clear, flat-bottom, 96-well plates (ThermoFisher). For the total serum antibody ELISA, these assay plates were coated with goat anti-mouse IgM or IgG1 polyclonal antibodies (Southern Biotech). Coating was done overnight at 4°C, with 100 µL of the antibody solution added to each well at a concentration of 3 µg/ml in PBS at pH 8.0. Subsequently, the plates were washed four times with PBST (0.1% Tween-20 in 1X PBS with Ca^2+^ and Mg^2+^) and blocked with 250 µL of ELISA diluent (eBioscience) for 3 hours at room temperature. After three times of washing with PBST, each well received 100 µL of serum samples or standards and was incubated at 4°C overnight. Both serum samples and standards were appropriately diluted with ELISA diluent. For serum samples, an 8-step 2-fold serial dilution was executed, commencing with a 1:8000 starting dilution. For standards, a 7-step 2-fold serial dilution was carried out, starting at a concentration of 40 ng/mL. Additionally, a blank well was included. Purified mouse IgM or IgG1 monoclonal antibodies (clones 11E10 or 15H6, respectively; Southern Biotech) served as standards. Technical duplicates were included for each sample. Following six washes with PBST, HRP-conjugated secondary antibodies specific for IgM or IgG1 (Southern Biotech) were diluted with ELISA diluent at a ratio of 1:2000 and incubated for 1.5 hours at room temperature in the dark (100 µL per well). After seven washes with PBST, 100 μL TMB substrate (eBioscience) was added to each well and developed for 3 minutes. The reaction was terminated by adding 100μL 1M phosphoric acid. Finally, the plates were read at 450 nm using a BioTek Synergy HT detector. Absolute concentrations of serum antibodies were determined by interpolation from the standard curve.

In the NP-specific ELISA, the first two columns on each plate were reserved for assay standards. These wells were coated with goat anti-mouse IgM or IgG1 polyclonal antibodies. After the coating step, these wells were blocked and subsequently incubated with different concentrations of purified mouse IgM or IgG1 monoclonal antibodies. These standards served as a reference for quantification. Remaining wells on the plate were coated with NP_(7)_-BSA or NP_(36)_-BSA (Biosearch Technologies) at a concentration of 3 µg/ml dissolved in 1X PBS pH 8.0. The wells were subsequently blocked and serum samples at varying dilutions were added to each well. The subsequent steps, including washing and detection, were similar to those used in the total serum antibody ELISA. Relative titers of NP-specific antibodies were determined by interpolating the sample absorbance values on the plate reference curve generated for each plate. This reference curve was established using purified mouse monoclonal IgM or IgG1 antibodies.

### STAT3 CUT&RUN

Naïve splenic B cells were initially cultured with αCD40 + IL-4 for 96 hours and IgM^+^ and IgG1^+^ B cells were sorted. 400,000 IgM^+^ or IgG1^+^ B cells were stimulated with αCD40 (1 μg/mL) + IL-4 (12.5 ng/mL) or αCD40 + IL-21 (10 ng/mL) for an hour, washed thrice with PBS, and resuspended in Antibody Buffer (1X eBioscience Perm/Wash Buffer, 1X Roche cOmplete EDTA-free Protease Inhibitor, 0.5 μM Spermidine, 2 μM EDTA in H_2_O). Cells were then incubated overnight with anti-STAT3 Mouse mAb (124H6, CellSignaling, Cat #9139, dilution 1/100) in Antibody Buffer in a 96 well V-bottom plate at 4°C. Cells were then washed twice with Buffer 1 (1X eBioscience Perm/Wash Buffer, 1X Roche cOmplete EDTA-free Protease Inhibitor, 0.5 μM Spermidine in H2O) and resuspended in 50ul of Buffer 1 + 1X pA/G-MNase (Cell Signaling, cat. 57813) and incubated on ice for 1 hour and washed thrice with Buffer 2 (0.05% w/v Saponin, 1X Roche cOmplete EDTA-free Protease Inhibitor, 0.5 μM Spermidine in 1X PBS). After washing, cells were resuspended in calcium Buffer (Buffer 2 + 2 μM CaCl_2_) for 30 mins on ice to activate the pA/G-MNase reaction, and equal volume of 2X STOP Buffer (Buffer 2 + 20 μM EDTA + 4 μM EGTA) was added along with 1 pg of Saccharomyces cerevisiae spike-in DNA (Cell Signaling, cat. 29987). Samples were incubated for 15 mins at 37°C and DNA was isolated and purified using Qiagen MinElute Kit according to manufacturer’s protocol and subjected to library amplification.

DNA was quantified by PicoGreen and the size was evaluated by Agilent BioAnalyzer. Illumina sequencing libraries were prepared using the KAPA HTP Library Preparation Kit (KAPA Biosystems #KK8234) according to the manufacturer’s instructors with <0.001-3.8 ng input DNA and 14 cycles of PCR. Barcoded libraries were run on the NovaSeq 6000 in a PE100 run, using the NovaSeq 6000 S4 Reagent Kit (200 Cycles) (Illumina). An average of 26 million paired reads were generated per sample.

Paired reads were trimmed for adaptors and removal of low-quality reads by using Trimmomatic^58^ (v0.39) and aligned to the mm10 reference genome using Bowtie2^59^ (v2.4.1) with --dovetail option. Upon alignment, peaks were called using MACS2^60^ (v2.2.7.1) with input samples as control using narrow peak parameters --cutoff-analysis -p 1e-5 --keep-dup all -B --SPMR. Tracks shown are bigwig files processed with deepTools^61^ (v3.5.1) bamCoverage with --binSize 10. Saccharomyces cerevisiae spike-in DNA scaled bigwig tracks from replicates were then averaged per condition using bigwigAverage function from deepTools with bin size of 10bp.

### Statistical analysis

Statistical analyses were conducted using GraphPad Prism 10 (GraphPad Software, Inc.). Graphs were generated using GraphPad Prism 10, and error bars indicate the mean with standard error. The specific statistical tests used for each analysis are detailed in the figure legends.

## ACKNOWLEDGEMENTS

We thank members of Chaudhuri lab for discussion and insightful feedback. J.C. was supported by grants from the National Institutes of Health (R01AI072194, R01AI124186 and P30CA008748), the Starr Cancer Research Foundation, the Ludwig Center for Cancer Immunotherapy, MSKCC Functional Genomics, and the Geoffrey Beene Cancer Center. We acknowledge the use of the Integrated Genomics Operation Core, funded by the NCI Cancer Center Support Grant (CCSG, P30 CA08748), Cycle for Survival, and the Marie-Josée and Henry R. Kravis Center for Molecular Oncology. We thank A. Bravo for help with maintenance of the mouse colony.

## AUTHOR CONTRIBUTIONS

Conceptualization, Y.K. and J.C.; Methodology, Y.K., F.M., S.G., W.T.Y., J.C.S., and J.C.; Validation, Y.K., F.M., S.G., and K.R.; Formal Analysis, Y.K., F.M., S.G., K.T.B., K.R., and; Investigation, Y.K., F.M., S.D.C., W.T.Y., and J.C.; Writing – Original Draft, Y.K. and J.C.; Writing – Review & Editing, Y.K., W.T.Y., and J.C.; Visualization, Y.K., K.T.B., and H.K.; Supervision, Y.K. and J.C.; Project Administration, Y.K., and J.C.; Funding Acquisition, S.D.C., and J.C.; Software, Y.K. and, K.T.B; Resources, S.G., and J.C.S.

