## Supplementary Figures for "IL-21 Shapes the B Cell Response in a Context-Dependent Manner"

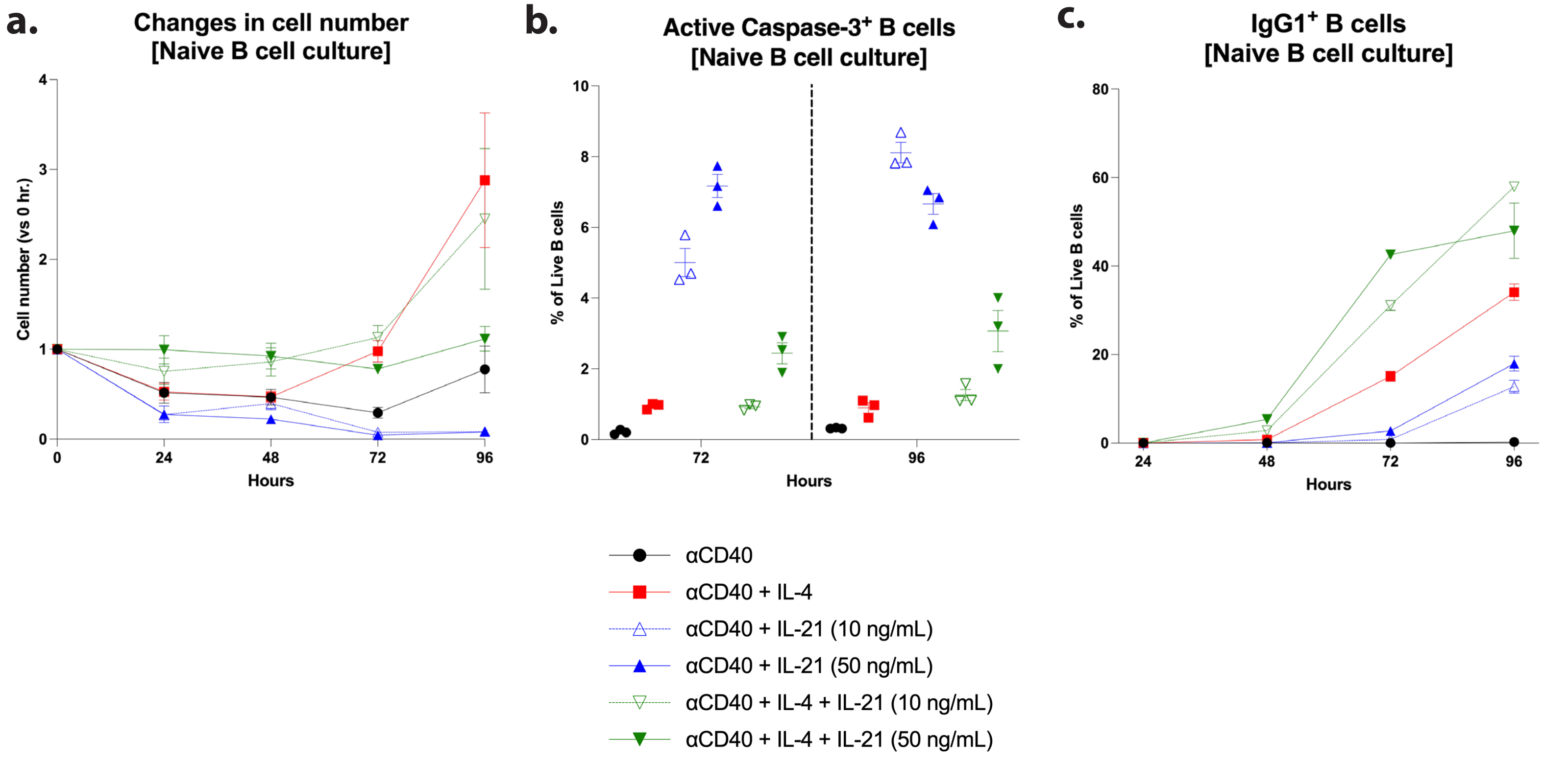


**Supplementary Figure 1**

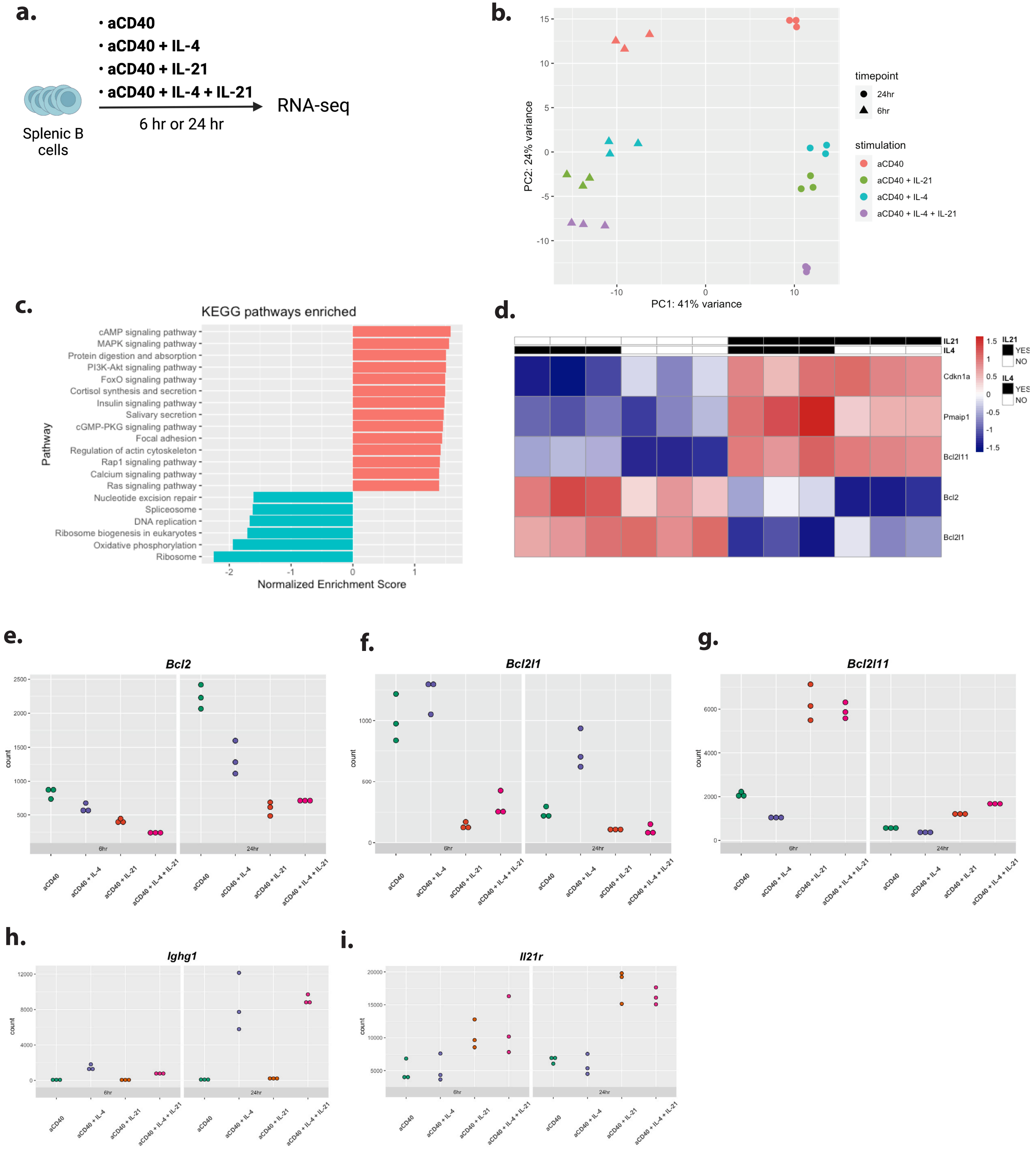


**Supplementary Figure 2**

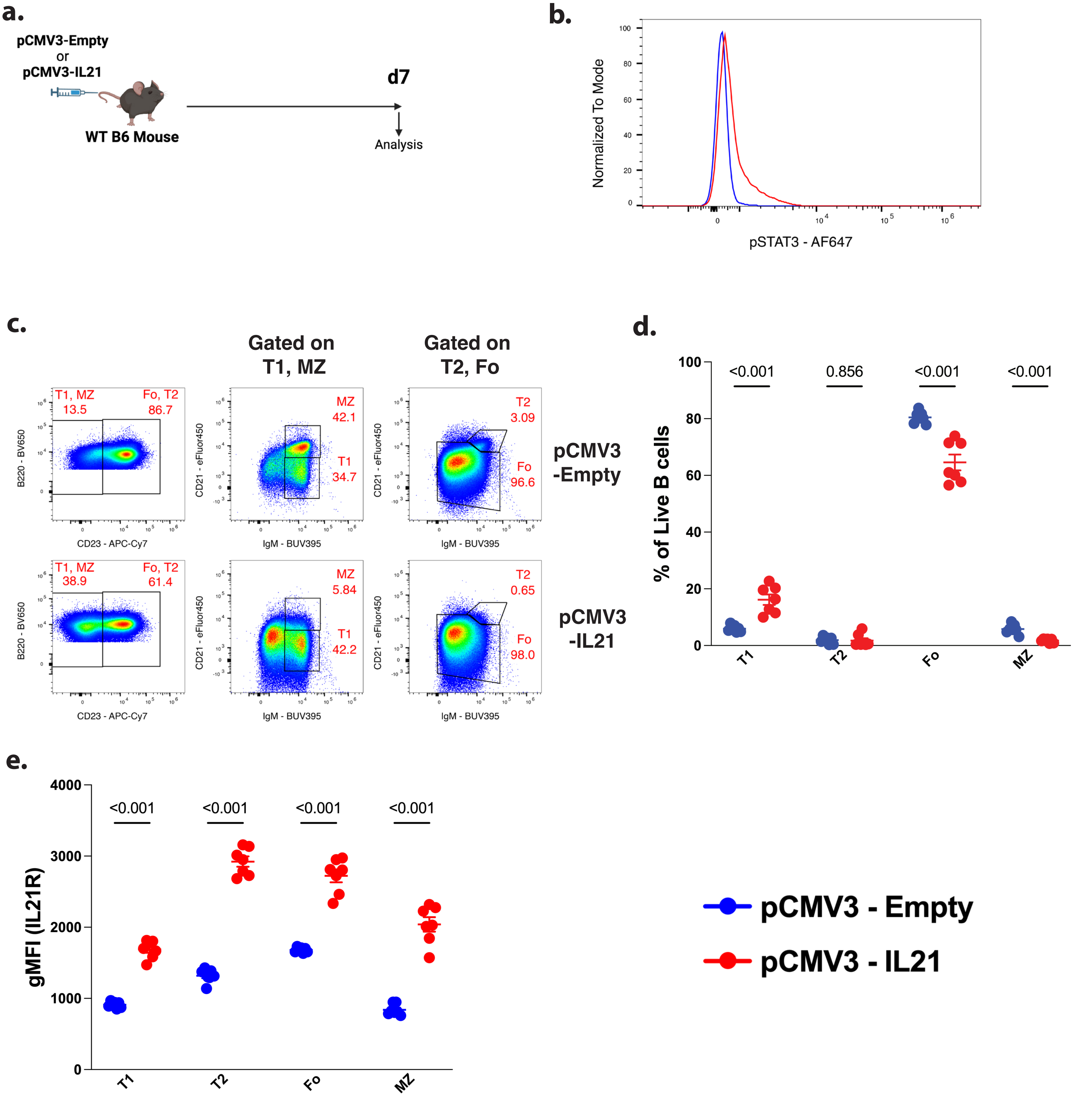


Supplementary Figure 3

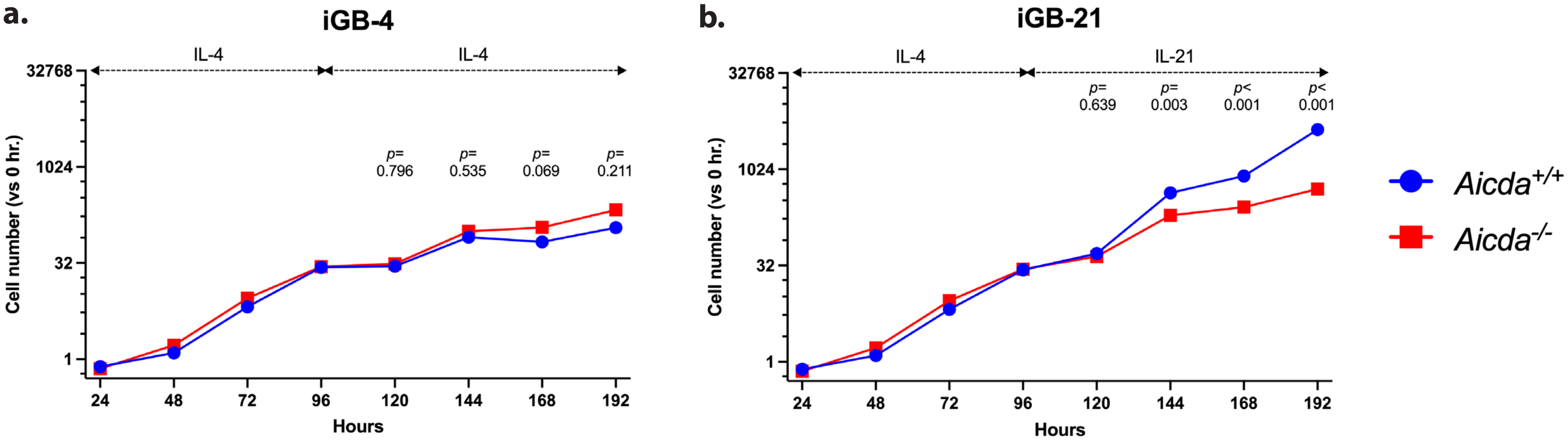


Supplementary Figure 4

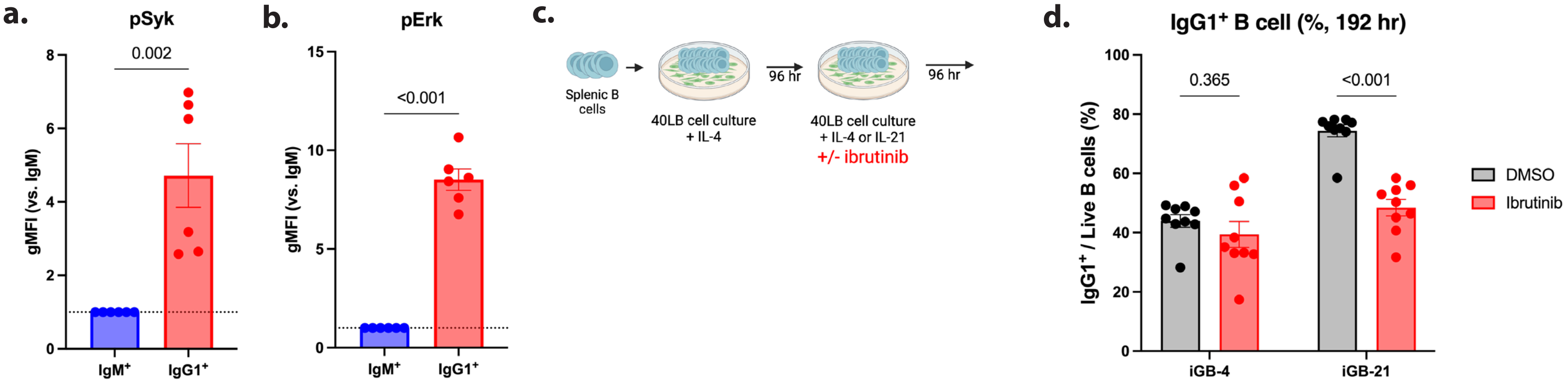


**Supplementary Figure 5**

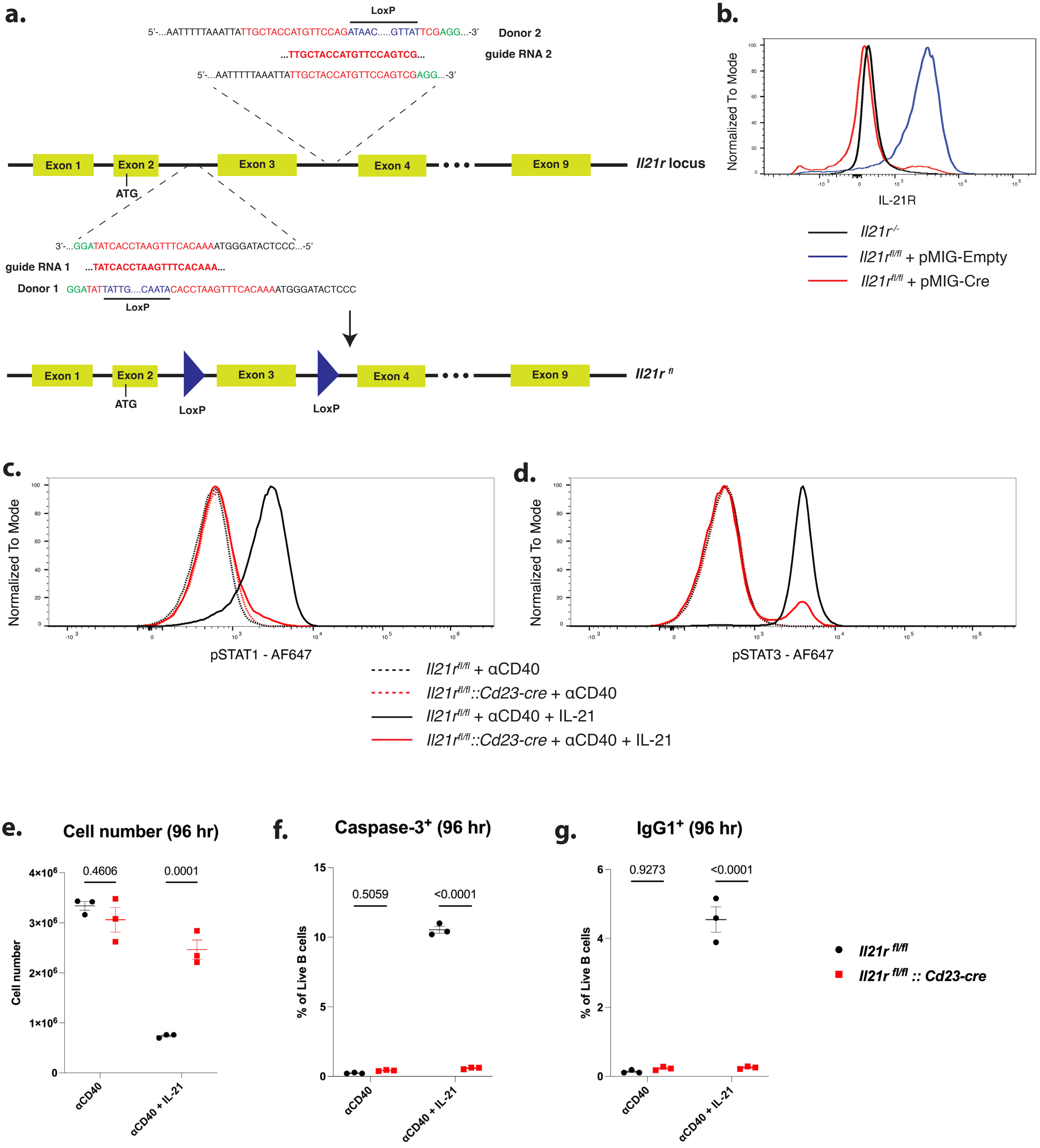


Supplementary Figure 6

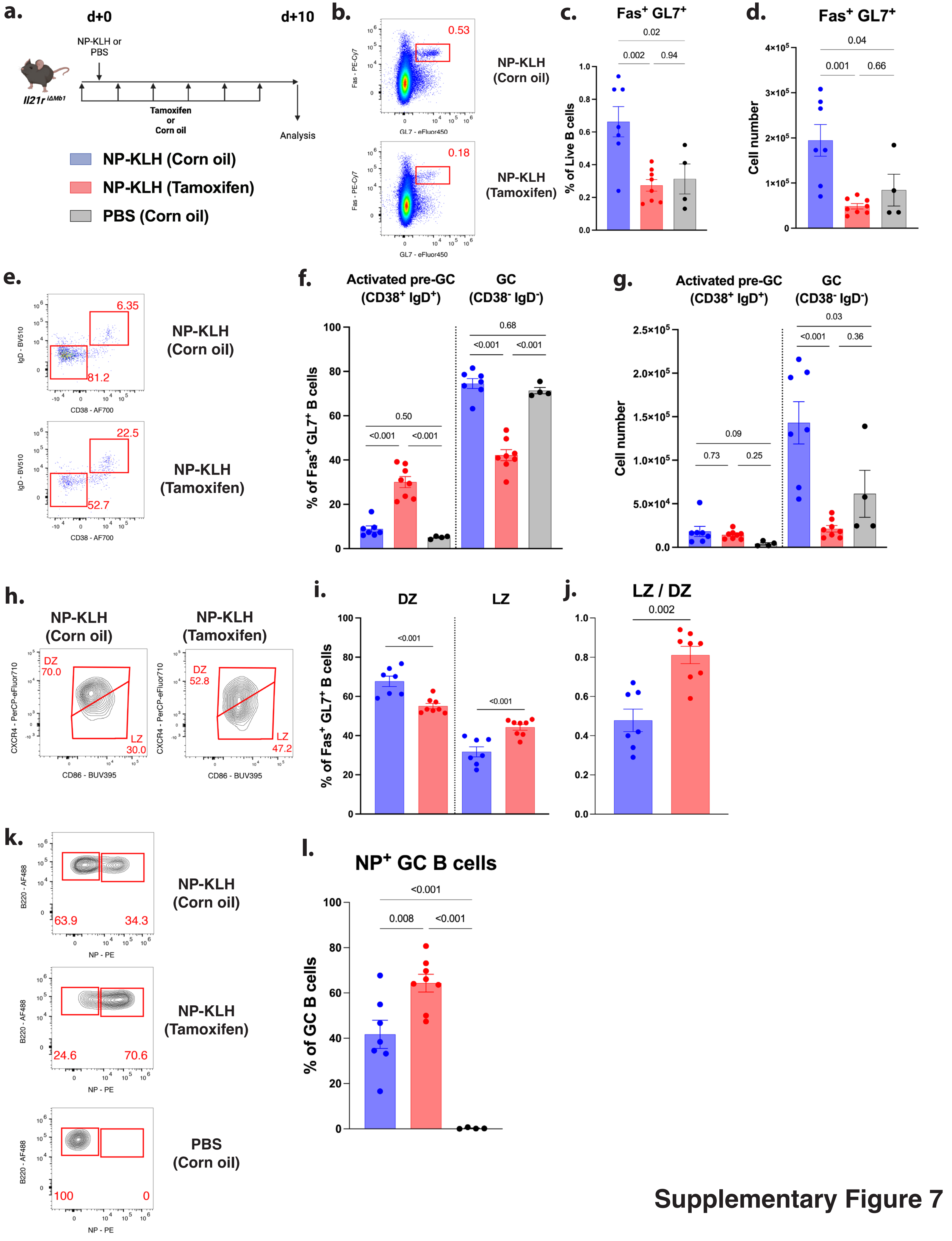


Supplementary Figure 7

### SUPPLEMENTARY FIGURE LEGENDS

**Supplementary Figure 1. Concentration-dependent effect of IL-21 on B cells *ex vivo*.**

(a) Changes in the cell count in culture over time. The y-axis represents changes in cell numbers relative to the starting point (0 hours). (b) Percentage of active caspase-3$^+$ B cells among total live B cells. (c) Percentage of IgG1-positive cells among naïve live B cells over the course of culture.

**Supplementary Figure 2. Transcriptomic changes in IL-21-stimulated B cells.**

(a) Experimental overview of the RNA-seq experiment. Naïve splenic B cells were cultured with αCD40 (1 μg/mL), IL-4 (12.5 ng/mL), or IL-21 (50 ng/mL) for 6 or 24 hours. (b) PCA plot demonstrating transcriptional differences among different stimulation conditions. Stimuli and timepoints are represented by distinct colors and shapes, respectively. (c) Differentially expressed KEGG pathway gene sets from MSigDB between αCD40 + IL-4 + IL-21 vs. αCD40 + IL-4-stimulated B cells (24-hour stimulation). Amongst pathways with an FDR < 0.05, pathways were prioritized based on the absolute value of normalized enrichment scores. A positive normalized enrichment score indicates enrichment in αCD40 + IL-4 + IL-21-stimulated B cells. (d) Heatmap illustrating the expression of pro-apoptotic and anti-apoptotic genes at the early (6-hour) timepoint. (e-i) Expression of (e) *Bcl2*, (f) *Bcl2l1*, (g) *Bcl2l11*, (h) *Ighg1*, and (i) *Il21r* under different stimulation conditions.

**Supplementary Figure 3. Transient *in vivo* overexpression of IL-21 leads to increased turnover of mature B cell population in spleen.**

(a) Experimental overview. (b) Representative histogram of phosphorylated STAT3 staining in the splenic B cell population. (c) Representative flow cytometry plot representing the gating strategy of follicular, transitional, and marginal zone B cell populations in the spleen. (d) Percentage of T1, T2, mature follicular (Fo), and marginal zone (MZ) B cells amongst splenic B cells. (e) Geometric MFI of surface IL-21R staining in T1, T2, Fo, and MZ B cells.

Different experimental groups are represented by distinct colors: pCMV3-Empty (blue) and pCMV3-IL-21 (red). Data were pooled from two independent experiments with *n* = 3 or 4 per experiment. Data in (d) and (e) were statistically analyzed with Student's unpaired t-test, with multiple comparisons correction applied using the Holm-Šídák method.

**Supplementary Figure 4. AID-deficient B cells proliferate less than AID-sufficient B cells in iGB-21, but not iGB-4 culture.**

(a, b) Changes in cell numbers of AID-sufficient or -deficient B cells over time in the (a) iGB-4 or (b) iGB-21 culture. The y-axis represents changes in cell numbers relative to the start of culture (0 hours). Data in (a) and (b) were statistically analyzed with Student's unpaired t-test, with multiple comparisons correction applied using the Holm-Šídák method.

**Supplementary Figure 5. Strong tonic BCR signaling in IgG1$^+$ B cells drives IL-21-mediated expansion of IgG1$^+$ B cells *ex vivo*.**

(a, b) Geometric MFI of (a) phosphorylated Syk and (b) phosphorylated Erk in the IgM$^+$ and IgG1$^+$ populations following 72 hours of αCD40 + IL-4 stimulation. The y-axis represents geometric MFI relative to the IgM$^+$ population. (c) Schematic representation of the iGB culture in the presence or absence of ibrutinib for inhibition of tonic BCR signaling. Naïve splenic B cells from WT mice were initially cultured on 40LB feeder cells and IL-4 (1 ng/mL) for 96 hours. Subsequently, cells were transferred to fresh 40LB feeder cells in the presence of IL-4 (1 ng/mL) or IL-21 (10 ng/mL), and either with DMSO or ibrutinib (100 nM). Feeder cells were replenished every 48 hours. (d) Percentage of IgG1$^+$ population among live CD19$^+$ B cells at 192 hours in the iGB-4 or iGB-21 culture. Data were pooled from two independent experiments with *n* = 3 - 5 per experiment. Data in (a), (b), and (d) were statistically analyzed using Student's unpaired t-test. For (d), multiple comparisons correction was applied using the Holm-Šídák method.

**Supplementary Figure 6. Generation and verification of the *Il21r-flox* mouse line.**

(a) Schematic representation of the *Il21r-flox* mouse line generation. The sequences of guide RNA to target the Cas9 nuclease to the upstream and downstream introns flanking exon 3 are depicted in red. The protospacer adjacent motif (PAM) is shown in green. The *LoxP* sites in the donor nucleotides are highlighted in blue. The resulting *Il21r*$^{fl}$ locus contains two *LoxP* sites flanking exon 3, and cre-mediated deletion of this exon leads to an early frameshift mutation. (b) Characterization of *Il21r-flox* mouse line using surface IL-21R staining. Naïve splenic B cells from *Il21r*$^{fl/fl}$ mice were cultured with αCD40 (1 µg/mL) + IL-4 (12.5 ng/mL) for 96 hours, with transduction at 48- and 72-hours using retrovirus carrying pMIG-Empty or pMIG-cre cassette for cre-mediated knock-out of IL-21R. Untransduced B cells from *Il21r*$^{-/-}$ mice were used as the control. *Il21r*$^{fl/fl}$ B cells transduced with pMIG-Empty and pMIG-cre, indicated in blue and red, respectively, were gated on GFP$^{+}$ (transduced) B cells, and untransduced *Il21r*$^{-/-}$ B cells, indicated in black, were gated on live B cells. (c, d) Representative flow cytometry plots of (c) phosphorylated STAT1 and (d) phosphorylated STAT3 after 30 minutes of stimulation with αCD40 (1 µg/mL) alone or αCD40 (1 µg/mL) + IL-21 (50 ng/mL) on naïve splenic B cells from *Il21r*$^{fl/fl}$ or *Il21r*$^{fl/fl}$*::Cd23-cre* mice. (e-g) (e) Cell number, (f) percentage of active caspase-3$^{+}$ cells, and (g) percentage of IgG1$^{+}$ cells after 96 hours of stimulation with αCD40 (1 µg/mL) alone or αCD40 (1 µg/mL) + IL-21 (50 ng/mL) on naïve splenic B cells from *Il21r*$^{fl/fl}$ or *Il21r*$^{fl/fl}$*::Cd23-cre* mice. Data in (e-g) were statistically analyzed with Student's unpaired t-test, with multiple comparisons correction applied using the Holm-Šídák method.

**Supplementary Figure 7. B cell-specific deletion of IL-21R leads to compromised GC responses.**

(a) Schematic representation of tamoxifen-induced B cell-specific knock-out of IL-21R in *Il21r*$^{i\Delta Mb1}$ mice and NP-KLH immunization. Mice were immunized with 100 µg NP-KLH precipitated in Imject Alum or 100 µL PBS. 3 mg of tamoxifen or 100 µL corn oil was administered to each mouse with oral gavage every two days, starting the 24 hours before immunization, for a total of 6 gavages.

Mice were euthanized and analyzed at day 10 post-immunization. (b) Representative flow cytometry plot representing $Fas^+$ $GL7^+$ B cell gating. The plot shown was pre-gated on live B cells. (c) Percentage of $Fas^+$ $GL7^+$ B cells among total splenic B cells. (d) Number of $Fas^+$ $GL7^+$ B cells per spleen. (e) Representative flow cytometry plot representing pre-GC ($IgD^+$ $CD38^{hi}$) and GC ($IgD^-$ $CD38^{lo}$) B cell gating. The plot was pre-gated on live $Fas^+$ $GL7^+$ B cell population. (f) Percentage of pre-GC and GC B cells among B $Fas^+$ $GL7^+$ B cells. (g) Number of pre-GC and GC B cells per spleen. (h) Representative flow cytometry plot representing DZ and LZ gating strategy. (i) Percentage of DZ ($CXCR4^{hi}$ $CD86^{lo}$) and LZ ($CXCR4^{lo}$ $CD86^{hi}$) B cells within live $Fas^+$ $GL7^+$ B cell population. (j) Ratio of LZ to DZ B cells within live $Fas^+$ $GL7^+$ B cell population. (k) Representative flow cytometry plot representing NP-stained GC B cells. The plot was pre-gated on live GC B cell population. (l) Percentage of NP-stained B cells within the GC B cell. Data were pooled from two independent experiments with $n = 3$ or 4 per experiment. Data in (c), (d), (f), (g), and (l) were statistically analyzed using ANOVA in conjunction with Tukey's multiple comparisons test. Data in (i) were statistically analyzed using Student's unpaired t-test. Data in (j) were statistically analyzed using Mann-Whitney unpaired non-parametric test.
